# Genetic encoding of 3-nitro-tyrosine reveals the impacts of 14-3-3 nitration on client binding and dephosphorylation

**DOI:** 10.1101/2022.11.11.516191

**Authors:** Phillip Zhu, Kyle T. Nguyen, Aidan B. Estelle, Nikolai N. Sluchanko, Ryan A. Mehl, Richard B. Cooley

**Affiliations:** Oregon State University, Department of Biochemistry and Biophysics, 2011 Agricultural and Life Sciences, Corvallis, OR 97331; A.N. Bach Institute of Biochemistry, Federal Research Center of Biotechnology of the Russian Academy of Sciences, 119071, Moscow, Russia

**Keywords:** nitration, 14-3-3, 3-nitrotyrosine, oxidative stress, phosphorylation, protein-protein interactions, genetic code expansion

## Abstract

14-3-3 proteins are central hub regulators of hundreds of phosphorylated “client” proteins. They are subject to over 60 post-translational modifications (PTMs), yet little is known how these PTMs alter 14-3-3 function and its ability to regulate downstream signaling pathways. An often neglected, but well documented 14-3-3 PTM found under physiological and immune-stimulatory conditions is the conversion of tyrosine to 3-nitro-tyrosine at several Tyr sites, two of which are located at sites considered important for 14-3-3 function: Y130 (β-isoform numbering) is located in the primary phospho-client peptide binding groove, while Y213 is found on a secondary binding site that engages with clients for full 14-3-3/client complex formation and client regulation. By genetically encoding 3-nitro-tyrosine, we sought to understand if nitration at Y130 and Y213 effectively modulated 14-3-3 structure, function, and client complexation. The 1.5 Å resolution crystal structure of 14-3-3 nitrated at Y130 showed the nitro group altered the conformation of key residues in the primary binding site, while functional studies confirmed client proteins failed to bind this variant of 14-3-3. But, in contrast to other client-binding deficient variants, it did not localize to the nucleus. The 1.9 Å resolution structure of 14-3-3 nitrated at Y213 revealed unusual flexibility of its C-terminal α-helix resulting in domain swapping, suggesting additional structural plasticity though its relevance is not clear as this nitrated form retained its ability to bind clients. Collectively, our data suggest nitration of 14-3-3 will alter downstream signaling systems, and if uncontrolled could result in global dysregulation of the 14-3-3 interactome.

## INTRODUCTION

Human 14-3-3 proteins (with isoforms denoted by their Greek letters β, γ, ε, ζ, η, θ, σ) exist as homo- and heterodimers and engage with hundreds of phosphorylated client proteins to regulate their catalytic activity, cellular localization, and ability to interact with other proteins.^1,2^ While 14-3-3 requires that client proteins be phosphorylated for complexation, 14-3-3 itself is subject to over 60 different post-translational modifications (PTMs)^1,3,4^ though little is understood about how these PTMs alter its function and the many essential signaling systems it controls. Among these PTMs is the conversion of several 14-3-3 tyrosine residues to 3-nitro-tyrosine (nY) in native, immune stimulated and oxidatively stressed tissues from rats, mice, and monkeys. ^5–10^ In contrast to most PTMs that are installed by “writer” proteins (e.g., kinases, acetyltransferases), tyrosine nitration occurs when superoxide (O_2_^-^) and nitric oxide (·NO) react to form the tyrosine nitrating agent peroxynitrite (ONOO^-^). ^11,12^ Interestingly, peroxynitrite reacts with only certain tyrosine residues on certain proteins,^6^ leading to the idea that protein nitration is not just an undesirable consequence of oxidative damage but can act as a regulated PTM imparting specific protein functional change that, like other PTMs when dysregulated, can lead to the onset of diseases. Given the importance of 14-3-3 in human physiology and disease, here we sought to uncover the structural and functional consequences of 14-3-3 nitration.

Proteomic mass spectrometry studies have revealed about half (but not all) of the tyrosine residues in 14-3-3 can be nitrated under physiologically relevant conditions, including immune system stimulation (Supporting Figure S1).^5–10^ Two of these, Y130 and Y213 (β-isoform numbering), are of particular interest because they are part of the primary and secondary client-interaction interfaces, respectively (Figure 1). The highly conserved primary site containing Y130 is the principle thermodynamic driver of client binding and is where the phospho-peptide of clients engage (Figure 1B). 14-3-3/client interactions at this interface are regulated by phosphorylation of client proteins at Ser/Thr sites flanked by specific recognition motifs denoted as Mode 1 (RXXp[S/T]XP), Mode 2 (RX[F/Y]Xp[S/T]XP) and C-terminally located Mode 3 (p[S/T]X_0-2_-COOH), though sequences that diverge from these canonical motifs have been established.^13^ These client phospho-motifs are usually found in unstructured or flexible regions^14^ enabling them to bind the deep and well conserved amphipathic groove of 14-3-3 containing the phospho-specific binding triad, of which Y130 is a part (Figure 1B). The secondary interface of 14-3-3/client interaction, of which Y213 is a part, is created by the C-terminal 3-helix bundle of 14-3-3 and is where domains of the client distal to the phospho-peptide engage with 14-3-3 (Figure 1C). This secondary engagement surface is well conserved among 14-3-3 isoforms. The number of structures of 14-3-3 bound to a full-length client are limited (Supplementary Figure S2)^15–23^, but in all eight the globular domain of the client docks with this surface patch and directly interacts with Y213. This interaction contributes to the overall thermodynamic stabilization of 14-3-3/client complexes,^21^ but also very important is that simultaneous client engagement at both primary and the secondary interfaces is required to achieve 14-3-3-based regulation of client activity, such as Raf kinase activation.^15,20^ Based on these observations, we hypothesized that nitration of 14-3-3 at sites Y130 and Y213 would alter its ability to interact with and regulate client function, and so *controlled* nitration at Y130 and Y213 could serve as a mechanism to regulate 14-3-3/client function, while *chronic* nitration of 14-3-3 could lead to dysregulation of phospho-protein signaling and disease.

**Figure 1.**
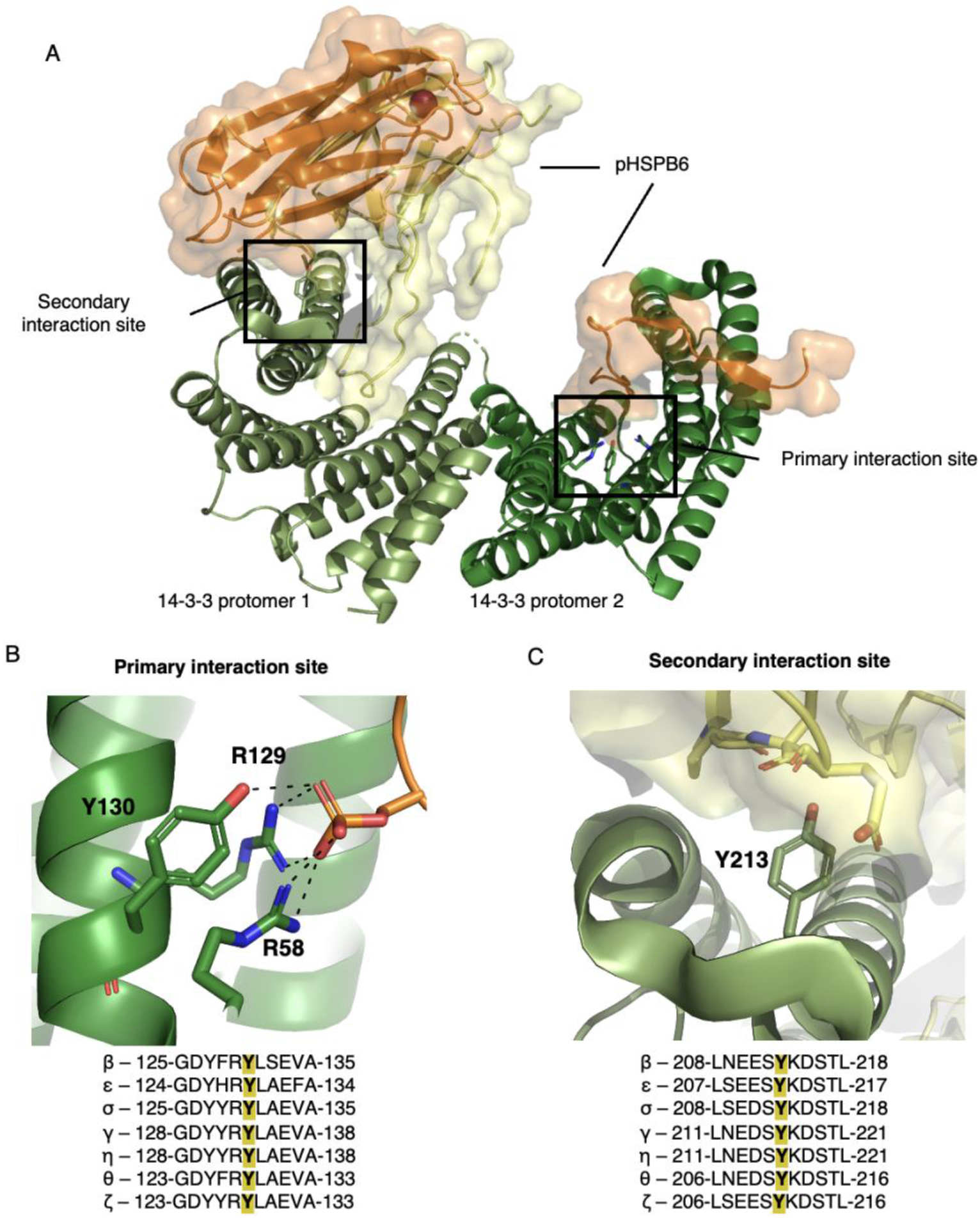
The general architecture of 14-3-3/client interactions. (A) The overall structural architecture of 14-3-3 (two protomers shown in pale green and dark green) when bound to the client pHSPB6 (orange and yellow cartoon with surface displayed, PDB ID: 5LTW), with major interactions highlighted in the secondary interaction site and the primary interaction site. (B) Zoom in of interactions formed between R58, R129, Y130 (β isoform numbering) with pHSPB6 phosphoserine (orange) in the primary interaction site. (C) Zoom in on the secondary interaction site of 14-3-3 with Y213 (green) shown interacting with pHSPB6 residues within 5 Å (yellow). Alignments of all 14-3-3 human isoforms, −5 and +5 residues of the tyrosine of interest (Y130 and Y213) are located under panels B and C, respectively.

Historically, studying the effects of tyrosine nitration has been challenging due to the use of chemical-based strategies to install nY in which proteins, cells or tissues are exposed to high concentrations of ^-^ONOO.^5,8,11,24–26^ These methods tend to oxidize many accessible tyrosine residues, as well as cysteines, methionines, and tryptophans, leaving studies on the site-specific effects of nitration out of reach.^27^ Highlighting this was elegant work by Smallwood et al. who showed that when calmodulin was exposed to ONOO^-^, over 200 unique PTM protein variants were produced.^28^ Indeed, the heterogeneity of peroxynitrite treated samples obfuscates functional assignment of any one specific nY PTM variant, and in part explains why to date only 4 different proteins with nY modifications at biologically relevant sites have been reported. ^29–32^

Genetic code expansion (GCE), on the other hand, permits site-specific incorporation of nY into proteins at genetically programmable amber (UAG) stop codons during translation using an orthogonal amino acid tRNA synthetase (RS)/tRNA pair.^33–39^ In order to study how nitration effects 14-3-3/client interactions, also needed is the ability to produce site-specifically phosphorylated client proteins, and for that here we also employ GCE systems to site-specifically install phosphoserine into a known client, Small Heat Shock Protein B6 (HSPB6).^21^ With these tools we solved the first crystal structures of PTM variants of 14-3-3 and by using *in vitro* binding assays show how nitration impacts phospho-client interaction. Specifically, we find that 14-3-3 nitrated at Y130, but not Y213, prevents complexation with phosphorylated-client proteins and increases their susceptibility to phosphatases. Lastly, using a mammalian nY GCE expression system, we evaluate the cellular localization of nitrated 14-3-3 and show that, unlike other client-binding deficient variants of 14-3-3 that typically localize to the nucleus, 14-3-3 nitrated at Y130 remains largely cytosolic. Collectively, our findings provide the first evidence that nitration of 14-3-3 influences its function and provide an example of how crosstalk between PTMs (nitration and phosphorylation) could work to rewire key signaling pathways.

## RESULTS

### Expression of site-specifically nitrated 14-3-3 β

We chose here to use the β-isoform due to its robustness of expression in our hands and its ability to be crystallized in both apo and peptide-bound forms.^40^ Residues Y130 and Y213 are strictly conserved across the 14-3-3 family and so insights into nitration of the β - isoform should be applicable to all isoforms. To express site-specifically nitrated 14-3-3 β, we introduced TAG codons at sites Y130 and Y213 and then used our recently developed *E. coli* nY GCE amber suppression system^33,34^ to incorporate nY at these sites to produce 14-3-3 β 130nY and 14-3-3 β 213nY, respectively. Expression of 14-3-3 β 130nY routinely produced protein at yields of ~ 50 mg per liter culture that by mass spectrometry was found to be homogenously nitrated (Supplementary Figure S3). On the other hand, expression of the 213nY form sometimes resulted in protein containing aminoTyr instead of nY (data not shown). The nY GCE system is not permissive for direct aminoTyr translational incorporation,^33,38^ and so we hypothesized this resulted from reduction of nY after translational installation. To overcome this issue, we found that expression of 14-3-3 β 213nY with a particularly higher rate of aeration (see Materials and Methods) was sufficient to produce homogenously nitrated protein (Supplementary Figure S3) at a similar yield to the 130nY form. That Y213 is fully solvent exposed while Y130 lies more buried in the amphipathic groove of 14-3-3 could explain why the former is more susceptible to reduction by cellular reductases or chemical reductants.

### Crystallization of 14-3-3 β nitrated at residues Y130 and Y213

We first sought to determine a molecular understanding for how site-specific nitration could impact 14-3-3 β structure using X-ray crystallography (Supplementary Table 1). 14-3-3 β 130nY crystallized readily in a condition and space group seen previously for 14-3-3 β,^40^ while 14-3-3 β 213nY grew in a new condition identified by sparse matrix screening. We discuss the individual structures in detail below.

### Nitration of Y130 alters the primary binding site of 14-3-3

The 1.5 Å resolution structure of 14-3-3 β 130nY showed an asymmetric structural topology similar to the previously determined apo 14-3-3 β wild-type structure (PDB code 2BQ0, 0.684 Å Cα RMSD), with one protomer adopting an ‘open’ conformation and the other the ‘closed’ form (Figure 2A, Supplementary Figure S4), as opposed to the more commonly observed, symmetrical doubly closed conformation.^22,40,41^ Unambiguous density for nY at site 130 was observed in both protomers (Figures 2B, C). In the open protomer (Figure 2B), the oxygen atom of the -NO_2_ moiety is placed just above the aromatic π-electrons of Y127 only 3.3 Å away. Overlays with the open protomer of wild-type 14-3-3 β structure show that the 130nY aromatic ring is rotated nearly 90° relative to that of Y130 in the wild-type structure (Figure 2D). Consequently, R58 is displaced such that its guanidinium group points outward and away from the amphipathic groove that comprises the primary binding site. The closed protomer resembles the conformational state of 14-3-3 β (and other isoforms) when bound to phospho-peptide, and so we overlayed this closed protomer with the structure of 14-3-3 β bound to the TFEB phospho-peptide to gain insight into whether 14-3-3 130nY would be able to bind clients (Figure 2E). This comparison showed that although the 130nY aromatic ring is positioned in a similar orientation as Y130 in the wild-type structure, the -NO_2_ moiety points toward the guanidinium groups of R58 and R129, pushing them away from the position they adopt when binding to phospho-peptide. We inferred these nitration-induced steric clashes would prevent the phospho-binding triad from adopting its binding-competent conformation, thereby preventing 14-3-3 from binding phosphorylated client proteins.

**Figure 2.**
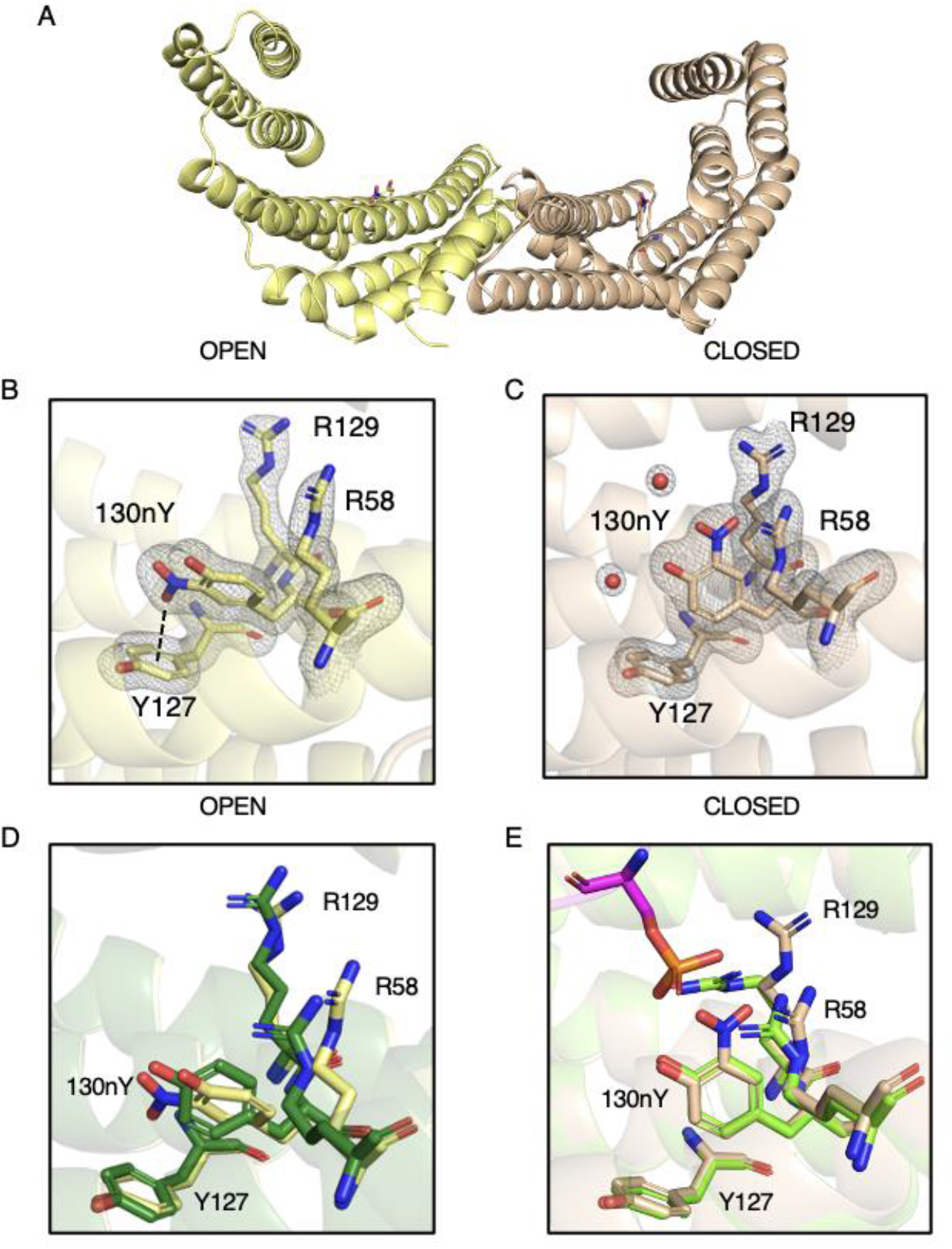
Structure of 14-3-3 β 130nY. (A) The biological dimer of 14-3-3 β 130nY shows one protomer (yellow) adopts an “open” conformation (yellow) and one a “closed” conformation (beige). (B) The structure of the phosphoserine binding triad (130nY, R129 and R58) as well as Y127 in the open protomer. Electron density for R58, Y127, R129, and 130nY and coordinating waters are shown. Dashed line represents the distance between -NO_2_ -oxygen to the center of Y127 aromatic ring. (C) The structure of the same phosphoserine binding triad in the closed protomer with its associated electron density is shown. All electron density (2Fo-Fc) is contoured to 1.5 σ. (D) An overlay of the open protomer phosphoserine binding site of 14-3-3 β 130nY (yellow) with the open protomer of wild-type apo 14-3-3 β (green, PDB ID: 2BQ0), (E) An overlay of the closed protomer phosphoserine binding site of 14-3-3 β 130nY (beige) with the closed protomer of wild-type apo 14-3-3 β bound to TFEB peptide (chartreuse, PDB ID: 6A5Q).

### Crystal structure of 14-3-3 nitrated at Y213 reveals an asymmetric domain-swap

Crystals of 14-3-3 β 213nY grew in different conditions and in a different space group than 14-3-3 β 130nY. In this 1.9 Å structure, both protomers adopt the ‘closed’ conformation (Supplementary Figure S5A), and densities for the nY residues are well defined (Supplementary Figure S5B, S5C). Each protomer of the dimer has a properly packed C-terminal ‘three helix bundle’ (helices α7, α8 and α9, residues 166 – 232), but interestingly tracing of the backbone electron density connecting helices α8 and α9 shows helix α9 is swapped with a neighboring lattice molecule in one of the protomers, but not both (Supplementary Figure S6). This assignment of asymmetric domain swapping is supported by B-factors analysis, which revealed the proposed domain-swapped protomer had markedly lower B-factors in both helices α8 and α9 (residues 185 – 232) than the protomer without density in this loop (Supplementary Figure S7). Lower B-factors for the domain swapped molecules would be expected since they have a more constrained packing interaction compared to those molecules not involved in domain swapping. While this domain swapping results in the formation of a tetramer (dimer of dimers), analytical size-exclusion chromatography of the 14-3-3 β proteins indicate both 130nY and 213nY were dimers in solution prior to crystallization, similar to 14-3-3 β wild-type (Supplementary Figure S8). The domain-swap therefore occurred during crystallization and may have been important for stabilizing crystal contacts. An analogous helix α9 domain swap was seen previously in a 14-3-3 ortholog from *Giardia duodenalis* (PDB: 4F7R), though in this case both protomers were engaged in the swap to form a filament like crystal lattice.^42^ What roles, if any, the formation of such domain-swapped multimers could play in 14-3-3 function is not clear. The fact that such a domain swap has not yet been observed before in a human 14-3-3 protein suggests Y213 nitration imparts an added level of structural pliability.

### Client binding to nitrated 14-3-3 β

We next tested whether nitration of 14-3-3 at sites Y130 and Y213 impair client binding. We first assessed the ability of 130nY and 213nY variants to bind R18, an engineered non-phosphorylated peptide that binds with high affinity (K_d_ ~70 nM for 14-3-3 β isoform) in the same amphipathic groove that phosphorylated peptides occupy.^43^ Since this peptide only binds to the primary binding site of 14-3-3, and the 130nY crystal structure indicated the key interacting residues R58 and R129 cannot adopt their binding competent conformations (Fig. 2, Fig. 3A), we would anticipate R18 should not bind the Y130 but should bind the Y213 nitrated variants. The R18 peptide was expressed in *E. coli* as a C-terminal fusion to sfGFP, while the 14-3-3 proteins were fused with an N-terminal AVI-SUMO tag (Supplementary Figure S9A). The AVI tag was used to install biotin at the N-terminus by BirA during co-expression for immobilization onto Streptavidin beads for pulldown assays (Supplementary Figure S9B), while SUMO served as an on-column proteolytic tag. After incubating 14-3-3 β wild-type, 130nY and 213nY proteins with sfGFP-R18 in solution, the 14-3-3 proteins were immobilized onto Streptavidin-Sepharose and any unbound sfGFP-R18 was washed away. Then, the 14-3-3 were eluted by SUMO protease (see Supplementary Materials and Methods) and any sfGFP-R18 that co-eluted was detected by SDS-PAGE analysis. These data revealed that sfGFP-R18 co-eluted with wild-type and 213nY variant, but not 130nY (Figure 3B), consistent with their respective structures.

**Figure 3.**
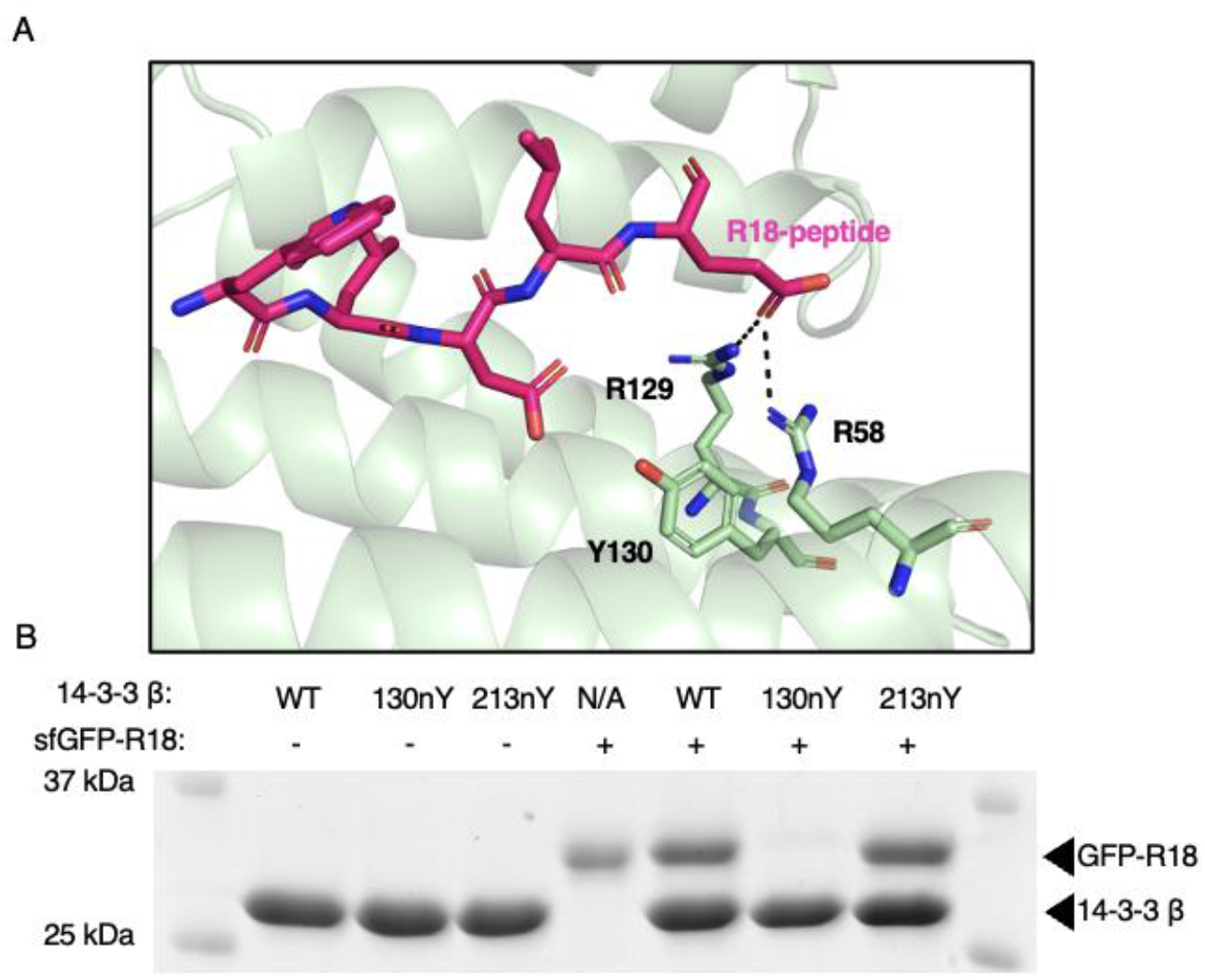
Assessment of R18 peptide binding to wild-type and nitrated 14-3-3 forms. (A) Crystal structure of engineered R-18 peptide (pink) bound to 14-3-3 (green) (PDB ID: 1A38) showing that this peptide binds the amphipathic groove where phosphorylated peptides and proteins bind. (B) SDS-PAGE analysis of pull-downs in which the indicated biotinylated SUMO-14-3-3 variants were first pre-incubated with and without sfGFP-R18, and then immobilized onto streptavidin beads. The beads were washed extensively and then 14-3-3 (along with any interacting sfGFP-R18 protein) was eluted with SUMO protease.

Since the R18 peptide is only expected to interact with the primary binding site of 14-3-3, we next sought to evaluate binding of nitrated 14-3-3 form to an authentic, biologically relevant phospho-protein in which Y130 directly engages with the phospho-group of the client and Y213 also engages with the bound client at the secondary interface. For this, we chose the well-studied client Small Heat Shock Protein B6 (HSPB6, also referred to as HSP20), which binds 14-3-3 when phosphorylated at Ser16 with low to mid-micromolar affinity.^21^ To generate HSPB6 phosphorylated at Ser16 (pHSPB6), we utilized the pSer3.1G system which combines the high efficiency pSer GCE chassis from Chin and colleagues with the healthy Release Factor 1 (RF1)-deficient expression host, B-95(DE3)Δ*A*Δ*fabR*Δ*serB*. ^44–46^ This system provides for homogenous incorporation of pSer into recombinantly expressed proteins, though hydrolysis of pSer can occur in the cell after incorporation. Because the expression host is RF1-deficient, little to no truncated protein is produced enabling the use of N-terminal affinity purification tags. SDS-PAGE of purified HSPB6 showed >95% purity, and Phos-tag gel electrophoresis confirmed the resulting pHSPB6 was ~80% phosphorylated (Figure 4, lanes 1 and 2).

**Figure 4.**
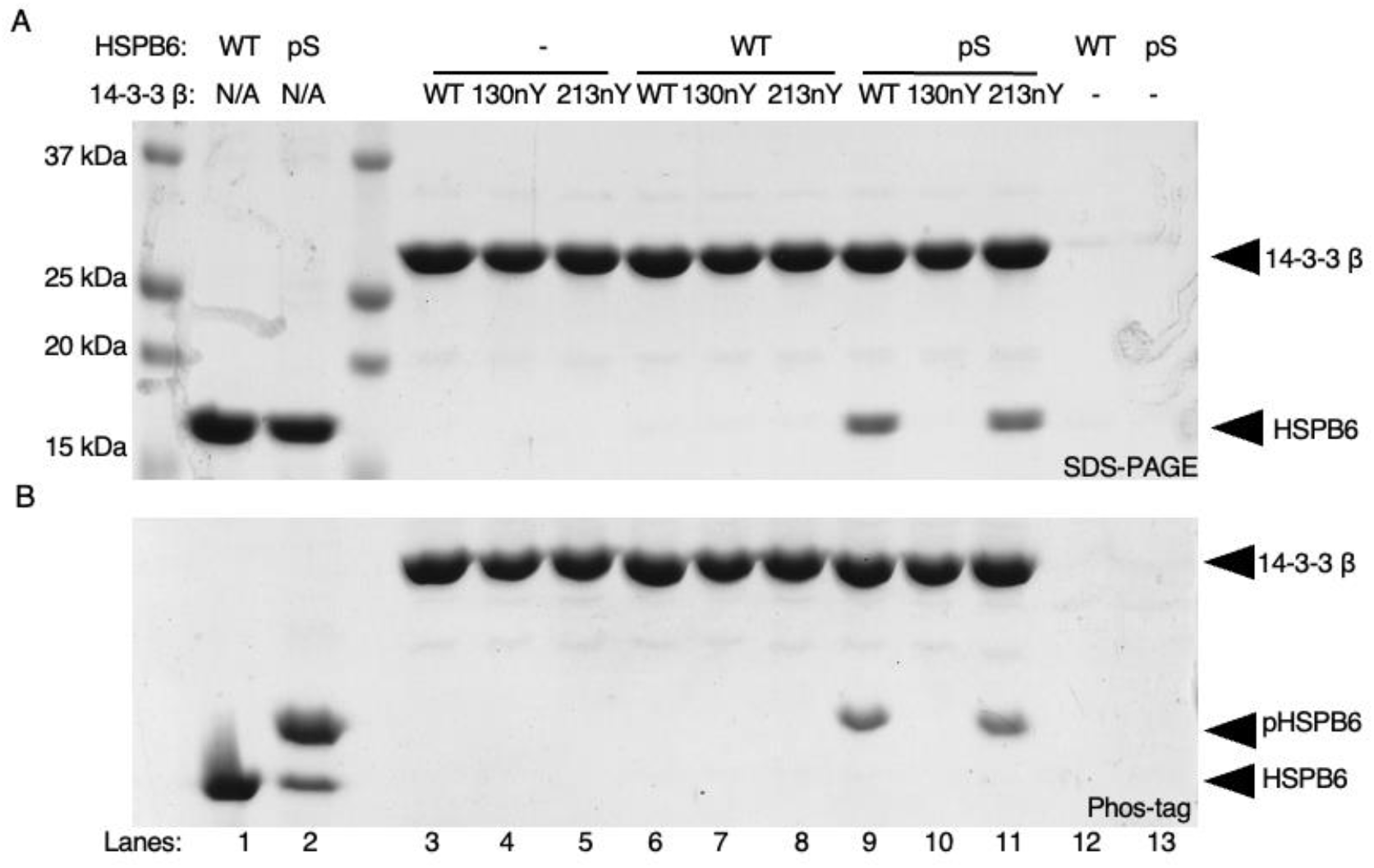
Interaction of nitrated 14-3-3 variants with full-length phosphorylated HSPB6. (A) SDS-PAGE and (B) Phos-tag gel analysis of purified HSPB6 (WT) and pHSPB6 (lanes 1 and 2), as well as pulldown assays in which biotinylated SUMO-14-3-3 β fusion proteins (WT, 130nY or 213nY) were incubated with no protein (lanes 3-5), HSPB6 (lanes 6-8) or pHSPB6 (lanes 9-11), immobilized onto Streptavidin beads and then eluted by SUMO protease after washing. Phos-tag electrophoresis has an acrylamide derivative that attenuates the migration of phosphorylated proteins, allowing for easy assessment of phosphorylation status of a protein.

The same biotinylated AVI-SUMO-14-3-3 β wild-type, 130nY and 213nY fusion proteins used for the R18 peptide pulldowns were incubated with wild-type HSPB6 and pHSPB6 to allow complexes to form. 14-3-3 β fusion proteins were again immobilized onto Streptavidin-Sepharose, and stable complexes were eluted after washing as done for the sfGFP-R18 pull down assay above. SDS-PAGE analysis revealed both wild-type and 14-3-3 β 213nY bound the pHSPB6 equally well (Figure 4). On the other hand, 14-3-3 β 130nY did not bind to pHSPB6. Phos-tag analysis of the eluted HSPB6 confirmed only the phosphorylated form bound to 14-3-3, further confirming that the interaction between 14-3-3 and HSPB6 was phosphorylation dependent (Figure 4B). Collectively, these data show that nitration of 14-3-3 β at Y130 – but not at Y213 – ablates its ability to stably bind phosphorylated client proteins. At the same time, these data do not exclude the possibility that Y213 nitration can affect the 14-3-3-regulated activity of some its clients.

In order to assess whether these results could be generalized to the broader 14-3-3 interactome, wild-type and nitrated 14-3-3 β proteins were immobilized onto NHS-Sepharose beads and incubated with the soluble fraction of a HEK293T cell lysate in which hundreds of phospho-clients are present.^2^ After incubation, the resins coupled with 14-3-3 β were collected, washed extensively, and interacting partners were released in denaturing buffer. SDS-PAGE analysis of the eluted protein pools was consistent with previous pull downs using R18 and HSPB6, showing that 130nY lacked the ability to stably bind client most all proteins pulled down while wild-type and 14-3-3 β 213nY were able to do so (Figure 5). The proteins that were pulled down with 14-3-3 β 130nY at ~30 KDa are likely endogenous 14-3-3 isoforms that heterodimerized with the immobilized protein. Indeed, the β-isoform of 14-3-3 is well established to heterodimerize with other isoforms of 14-3-3^47^ and our crystal structure confirmed the dimerization interface of 14-3-3 β 130nY is intact and unaltered relative to wild-type, and therefore should be able to form expected heterodimers. No proteins were pulled down with resin lacking 14-3-3 β protein, indicating the eluted proteins were dependent on interactions with 14-3-3 β.

**Figure 5.**
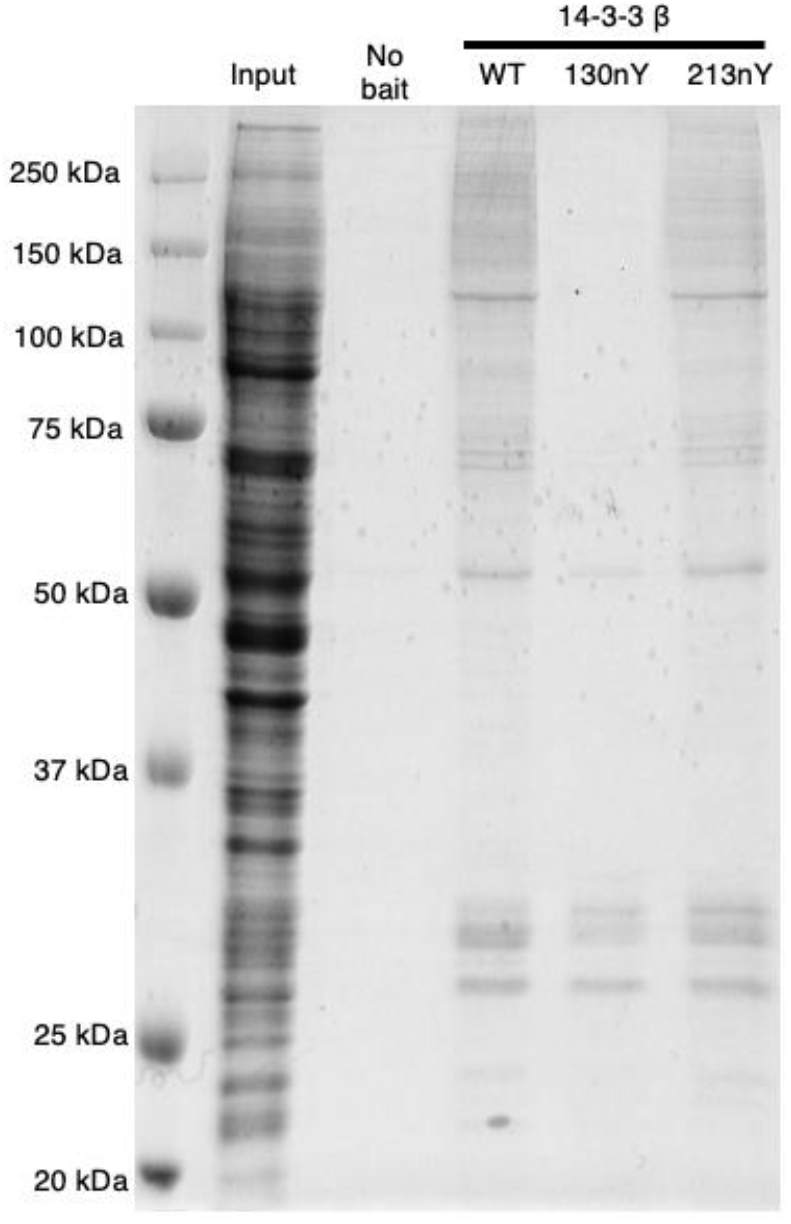
Analysis of nitrated 14-3-3 interactions with proteins from HEK293T cell lysate. Sepharose-conjugated 14-3-3 variants (or Sepharose conjugated to Tris; “No bait”) were incubated with HEK293T lysates, after which resins were collected, washed and then the interacting proteins were eluted with denaturant. Gel was stained with Coomassie Blue. “Input” lane is the HEK293T lysate prior to incubation with 14-3-3 variants.

### Dephosphorylation of pHSPB6

14-3-3 is well known to protect clients from dephosphorylation by phosphatases.^48^ Based on the above experiments, we reasoned that nitration at Y130 could release clients thereby exposing them to dephosphorylation by phosphatases, whereas nitration at Y213 might not affect 14-3-3 mediated protection of clients from phosphatases. Using Phos-tag electrophoresis of pHSPB6, we evaluated the rate of pHSPB6 dephosphorylation by catalytic amounts of λ phosphatase (~1 enzyme : 2000 substrate molar ratio) in the presence of different nitrated 14-3-3 β variants (Figure 6). The addition of phosphatase to pHSPB6 in the absence of 14-3-3 led to the majority being dephosphorylated within 30 min, and complete dephosphorylation after 3 hours (Figure 6 A,B). When pHSPB6 was pre-complexed with 14-3-3 β wild-type, only ~ 30% was dephosphorylated after 3 hours, confirming the protective effects of 14-3-3 β on client phosphorylation status (Figure 6 C,D). The 14-3-3 β 213nY variant afforded a similar level of protection for pHSPB6 against dephosphorylation as 14-3-3 β wild-type. In contrast, the 14-3-3 β 130nY provided little to no protective effect with all pHSPB6 being dephosphorylated after 3 hours (Figure 6 A,C), consistent with it not being complexed to 14-3-3. Collectively, these data indicate that nitration at site Y130, but not Y213, promotes the dissociation of clients and increases their susceptibility to dephosphorylation by phosphatases.

**Figure 6.**
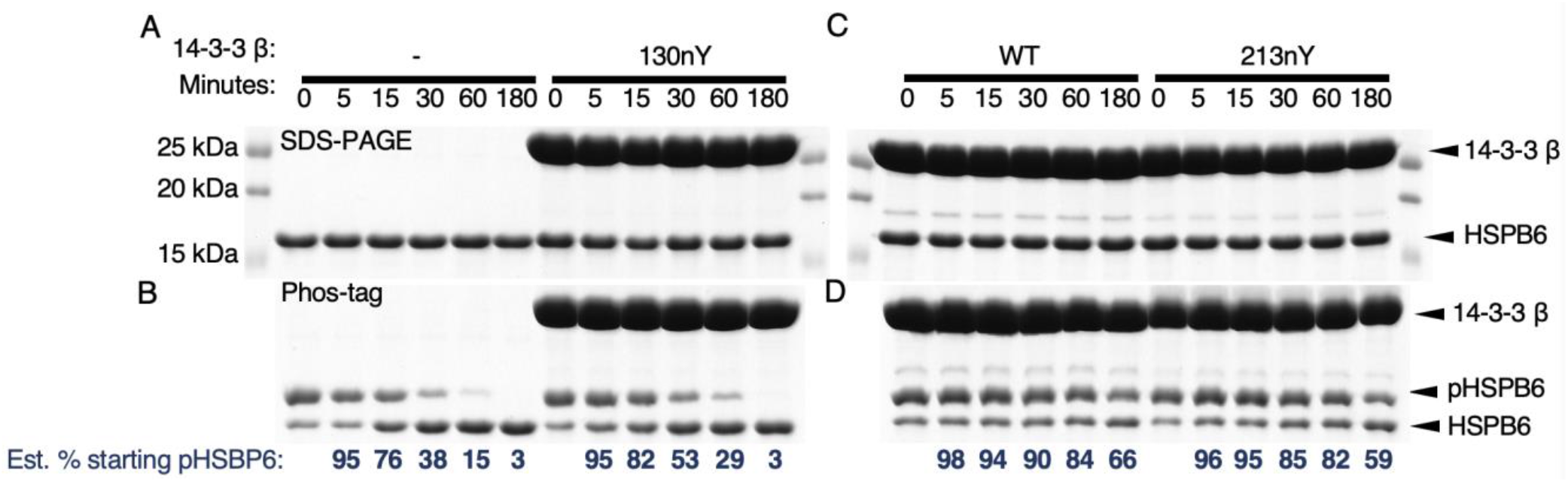
Dephosphorylation of pHSPB6 in the presence of nitrated 14-3-3 β. pHSPB6 was pre-incubated with buffer only, 14-3-3 β WT, 130nY, or 213nY and then subjected to dephosphorylation by λ phosphatase (see Materials and Methods). Samples were taken at 0, 5, 15. 30, 60, and 180 minutes for analysis by SDS-PAGE (A, C) or Phos-tag gel electrophoresis (B, D). Relative percentage of the starting amount of pHSPB6 were estimated by densitometry and shown below each Phos-tag gel.

### Localization of nitrated 14-3-3 β in mammalian cells

Previously, it has been demonstrated that wild-type 14-3-3 is largely localized to the cytoplasm, while client-binding deficient variants with mutations in residues involved in phospho-amino acid interactions (e.g., K51E, β numbering) localize to the nucleus.^49–54^ Having shown here that nitration of 14-3-3 β at position Y130 greatly attenuates its affinity to clients *in vitro* by compromising phospho-amino acid interactions, we asked if 14-3-3β 130nY would also localize to the nucleus. To do this, HEK293T cells were co-transfected with our mammalian nY GCE machinery^35^ and plasmids expressing either wild-type (with and without a nuclear localization sequence [NLS]) or the K51E, 130TAG and 213TAG 14-3-3 β variants fused to eGFP at the C-terminus. The fused eGFP was added as a fluorescence tag to track cellular localization, and, because it is located at the C-terminus of 14-3-3, only protein in which the TAG codon was successfully suppressed by nY incorporation would be visualized. Total fluorescence produced by the cells were similar for wild-type, NLS-wild-type, and the K51E mutants of 14-3-3 β-eGFP fusion proteins, while Y130TAG and Y213TAG mutants expressed at ~66% and ~30% that of wild-type, consistent with the lower translational efficiency of the GCE TAG codon suppression system (Supplementary Figures S10 and S11). Importantly, no fluorescence was observed in the TAG mutant expressing cells when nY was omitted from the media, confirming accurate incorporation of nY (Supplementary Figure S10 and S11). Fluorescence imaging of fixed cells after expression confirmed wild-type 14-3-3 β-eGFP protein localized predominantly to the cytoplasm, while the NLS-wild-type and K51E mutant 14-3-3 β proteins localized to the nucleus (Figure 7 A-F). Interestingly, 14-3-3 β 130nY was largely localized to the cytoplasm like wild-type 14-3-3 β (Figure 7 G,H), even though nitration at Y130 inhibits client binding and therefore should lead to nuclear localization, like K51E (Figure 7 G, H). The 213nY 14-3-3 β-eGFP also localized to the cytoplasm like wild-type and 130nY (Figure 7 I, J). Collectively, these data demonstrate that the client-binding defective 14-3-3 β 130nY does not traffic to the nucleus like other binding defective mutants, and that there may be additional regulatory factors at play that determine the function and fate of nitrated 14-3-3.

**Figure 7.**
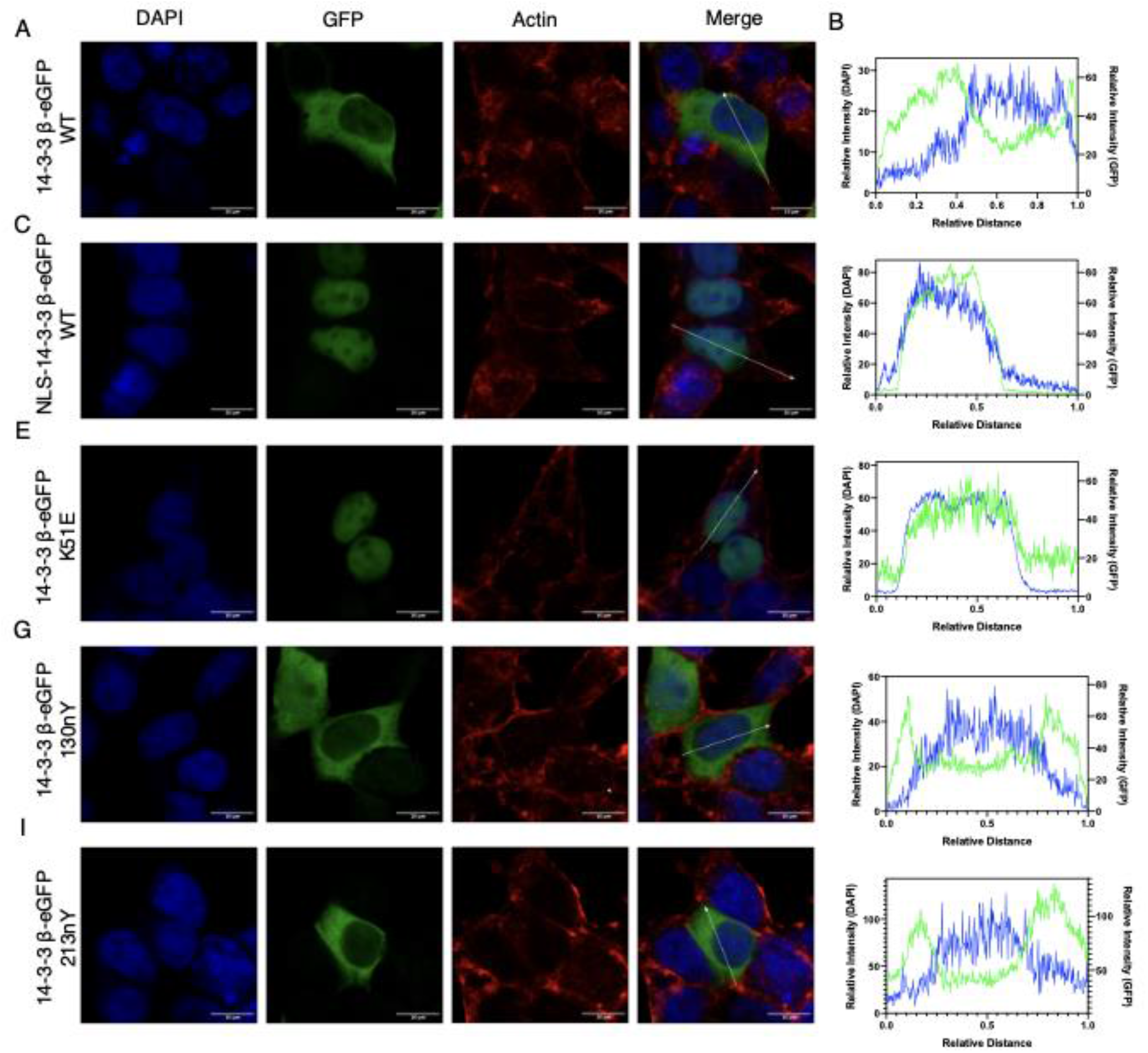
Subcellular localization of nitrated 14-3-3. HEK293T cells expressing 14-3-3 β-eGFP fusion proteins (green channel) were stained for the nucleus with DAPI (blue channel) and for actin with Alexa Flour 647 Phalloidin (red channel) and imaged by confocal microscopy. Subcellular fluorescence profile quantification of DAPI and eGFP are shown in far-right panels. (A,B) 14-3-3 β-eGFP WT, (C,D) NLS-14-3-3 β-eGFP WT, (E,F) 14-3-3 β-eGFP K51E, (G,H) 14-3-3 β-eGFP 130nY, and (I,J) 14-3-3 β-eGFP 213nY. See Supplementary Figure S9 for uncropped images.

## DISCUSSION

### PTMs on 14-3-3 as a mechanism of regulated client release

Over 60 PTMs have been mapped to 14-3-3, including phosphorylation, acetylation, mono- and di-methylation, ubiquitination and succinylation.^3^ How each PTM effects client binding – and whether these effects are 14-3-3 isoform and/or client-specific - remains an open question. To date, lysine acetylation at K49 and K120 (ζ-numbering - residues directly involved in phospho-peptide binding), as well as phosphorylation at S186 (β-numbering), T232 (ζ-numbering) were found to inhibit or release client-binding, while phosphorylation at S58 (ζ-numbering) monomerizes 14-3-3.^55–65^

Here, using (i) an engineered 14-3-3 binding peptide, (ii) an authentic full-length phosphorylated client, and (iii) the large pool of cellular clients from HEK293T cell lysates, we add to this short list of known 14-3-3 PTM effects by showing site-specific nitration of Y130 prevents clients binding. The crystal structure of 14-3-3 130nY revealed that by adding an -NO_2_ group to Y130, neighboring residues R58 and R129 are sterically hindered from adopting conformations required to form the functional “pSer binding triad”. We presume by extension that nitration at Y130 serves as a mechanism to release pre-bound clients, though directly testing nitration-induced release of clients is challenging, as *in vitro* nitration via bulk addition of peroxynitrite will oxidize a combination of tyrosines, cysteines, methionines and tryptophans,^28,66^ and delineating the contribution of each unique modification is not feasible. It is reasonable to expect that thermal fluctuations and the small size of peroxynitrite make it feasible Y130 could be nitrated when pre-bound to a client despite the relatively low solvent accessibility of this residue. With regards to nitration at Y213, we did not detect an obvious effect on client binding, but it cannot be ruled out that client-specific changes still occur given that proper 14-3-3 regulated client function (e.g. catalytic activity) requires engagement at both primary and secondary 14-3-3 interaction sites.^15,20^ In other words, while the 14-3-3/client complex may still be thermodynamically stable, 14-3-3-regulated client function may be altered by nitration at Y213. Further, the domain swapping of the C-terminal helix observed in the 213nY structure alludes to an added level of structural plasticity not yet observed in human 14-3-3 proteins. The only other 14-3-3 structure with a similar helix α9 domain swap was that from *G. duodenalis* (PDB ID: 4F7R), resulting in the formation of supramolecular assemblies that, interestingly, was controlled by phosphorylation at residue T214 – a residue in the loop connecting α8 and α9 and therefore a similar position as Y213 of human 14-3-3 β. Perhaps chronic nitration at Y213 could promote similar fibril like aggregates of human 14-3-3 in cells.

### Potential for 14-3-3 nitration affecting signaling systems

Given the role 14-3-3 plays as a central hub protein for hundreds of signaling systems, the functional changes imparted by tyrosine nitration could offer a mechanism to connect cellular oxidative stress with disease. As an initial foray into this, we leveraged here our recently developed GCE system for expressing nY proteins in mammalian cells,^35^ and first asked whether nitration of 14-3-3 altered its cellular localization. Prior work has shown that in the absence of bound clients, 14-3-3 will localize to the nucleus where it can recruit new clients and shuttle them back to the cytoplasm.^53^ To our surprise, 14-3-3 β 130nY was predominantly cytosolic even though it should not bind clients. This observation might be explained in part by the fact that 14-3-3 β 130nY expressed in mammalian cells will heterodimerize with native 14-3-3 molecules, making hemi-binding deficient 14-3-3 dimers (i.e., one protomer nitrated and one unmodified), though this is also expected when expressing 14-3-3 β K51E yet this variant still localizes predominantly to the nucleus. Localization of 14-3-3 to the nucleus therefore may not simply be an outcome of having released a client, rather such localization may be determined by the origin of the release, or other unknown secondary impacts caused by tyrosine nitration. Nevertheless, we expect that the inability of 14-3-3 β 130nY to bind clients will in and of itself alter 14-3-3 signaling systems. An important question to ask in this regard is, what proportion of 14-3-3 protein would need to be modified to elicit a physiological response? With 14-3-3 being one of the most abundant proteins in the cell (accounting for 0.1-1.3% of total soluble cellular proteins), clients released by 14-3-3 nitration can rebind another free, unmodified 14-3-3 molecule, or displace more weakly bound clients.^67^ But, free (unbound) phospho-clients are not protected from phosphatases as they are when bound to 14-3-3 (Figure 6) and so even transient release of clients could lead to their dephosphorylation and change the distribution of bound to unbound clients (Figure 8). In other words, low levels of 14-3-3 nitration could be sufficient to promote prolonged client release, dephosphorylation and therefore the controlled triggering of downstream signaling systems. In diseased cells, excessive (stoichiometric) nitration of 14-3-3 could lead to an uncontrolled release of clients, leading to the dysregulation of many signaling systems and a possible a connection between oxidative stress and disease. Nevertheless, given the plethora of PTMs found on 14-3-3, it will be critical to understand how they too alter the 14-3-3 interactome and the countless signaling systems under its control. In this regard, GCE technologies that encode not only nY but also other PTMs such as phosphoserine, phosphothreonine and acetyl-lysine will prove indispensable.

**Figure 8.**
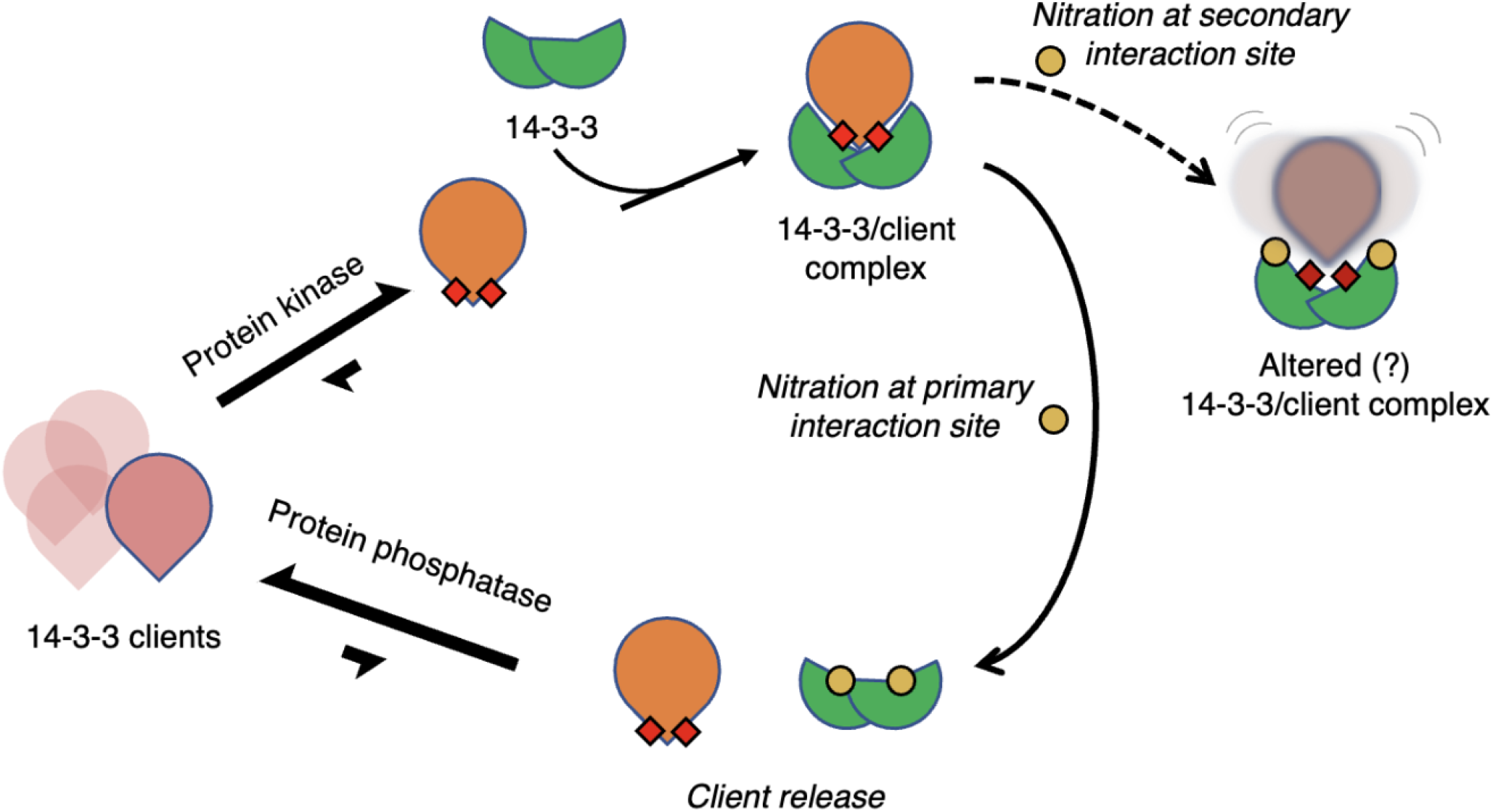
Schematic representation of the impact of 14-3-3 nitration and phospho-client binding. 14-3-3 (green) binds to 14-3-3 clients (pink – unphosphorylated, orange with red squares – phosphorylated) upon phosphorylation by protein kinases. Nitration of tyrosine (yellow dots) at the secondary interaction site may cause altered 14-3-3/client complexes while nitration of tyrosine at the primary interaction site promotes client release and increases client susceptibility to protein phosphatases.

## MATERIALS and METHODS

Below are brief descriptions of the Materials and Methods. Please see Supplementary Materials for more details.

### Bacterial cell lines

#### E. coli strains

BL21-ai (ThermoFisher, catalog no. C607003) and DH10B (ThermoFisher, catalog no. EC0113) were purchased from Thermofisher Scientific. B-95 Δ*A* Δ*fabR* Δ*serB* was created by the GCE4All Center (previously the Unnatural Protein Facility) as previously described.^44^ The PPY strain of *E. coli* used for generating SLiCE cloning extract was a generous gift from Y. Zhang (Albert Einstein University).

### Plasmid generation

#### Constructs for protein expression in E. coli

All construct assemblies for *E. coli* expression plasmids utilized the PPY-based SLiCE method and were transformed into DH10b cells.^68^ Genes encoding 14-3-3ζ, HSPB6, R18, *bd*SUMO^69^, *bd*SUMOEu1^70^, *bd*SENP1 and *bd*SENP1-EuB were codon optimized for *E. coli* expression and chemically synthesized (Integrated Technologies, Coralville, Iowa). The wild-type 14-3-3 ζ gene was cloned into NcoI/XhoI digested pBAD plasmid (Addgene #85482) with a TEV cleavable C-terminal His6 purification tag. Amber TAG codons were introduced at Y130 and Y213 using SLiCE. N-terminal AVI-tag^71^ and *bd*SUMOEu1 tags with short linkers between them were appended to 14-3-3 β for pull-down experiments. The *bd*SUMOEu1 tag is a variant of *bd*SUMO that is orthogonal (uncleavable) by eukaryotic SUMO proteases but can be cleaved by its orthogonal *bd*SENP1-EuB protease. For encoding 3-nitro-tyrosine, 14-3-3 β constructs were co-expressed with the pDule2-3NY-A7 machinery plasmid. For *in vivo* biotinylation, the AVI tagged 14-3-3ζ constructs were co-expressed with BirA from the pEVF plasmid as previously described.^72^ For *bd*SUMO and *bd*SUMO-Eu1 proteolytic cleavage, *bd*SENP1 and *bd*SENP1-EuB were expressed with a TEV cleavable N-terminal His6 tag from the pET28 vector, respectively.

To express HSPB6, a His6-*bd*SUMO-HSPB6 construct was cloned into NdeI/XhoI digested pRBC plasmid (Addgene #174076) and then a TAG codon was inserted at position S16 using SLiCE. These plasmids were co-expressed with the pKW2-EFSep phosphoserine machinery plasmid (Addgene #173897). All amino acid sequences of recombinantly expressed 14-3-3 constructs can be found in the Supplementary information provided.

#### Constructs for protein expression in E. coli

For incorporating 3-nitro-tyrosine into proteins in HEK293T cells, the pAcBac1 plasmids expressing 14-3-3ζ WT, Y130TAG and Y213TAG fused C-terminally with sfGFP-His6 were generated as previously described.^35^ An N-terminal NLS sequence and the K51E mutation were added to the wild-type construct by overlap-extension PCR, and then ligated into NheI/EcoRI digested pAcBac1 plasmid. The 3-nitro-tyrosine incorporation plasmid (pAcBac1-3-nitroTyr-A7, Addgene # 141173) was as previously described.^35^

### *E. coli* Protein Expression and Purification

All 14-3-3ζ variants and sfGFP-R18 were expressed in *E. coli* BL21-ai cells using auto-induction media at 25 °C, purified with standard metal affinity methods and eluted with the addition of buffer containing 300 mM imidazole. 3-nitro-tyrosine and D-biotin were supplemented to the media at 0.5 mM and 100 μM, respectively, when needed. Wild-type HSPB6 and HSPB6 with phosphoserine at site S16 were expressed in *E. coli* B95(DE3)Δ*A* Δ*fabR* Δ*serB* in auto-induction media at 25 °C, purified with standard metal affinity methods. *bd*SENP1 and *bd*SENP1-Eu1 proteases were expressed in BL21ai cells in 2xYT media at 18°C with 1 mM IPTG/0.1% (w/v) arabinose for 18 hrs. 14-3-3ζ and HSPB6 proteins were further purified by size-exclusion chromatography as needed, concentrated, and frozen at −80 °C. Further details can be found in the Supplementary Materials.

### Mass Spectrometry

#### Whole protein mass spectrometry

14-3-3ζ wild-type, 130nY and 213nY proteins were buffer exchanged into 50 mM ammonium acetate, concentrated using EMD Millipore C4 resin ZipTips, and analyzed using an FT LTQ mass spectrometer at the Mass Spectrometry Facility at Oregon State University.

### Protein Crystallization and X-ray crystallography

#### Protein Crystallization, Data Collection and Processing

Crystals were grown by the hanging drop method in which 14-3-3ζ 130nY and 213 nY proteins at 20-30 mg/ml in 50 mM HEPES pH 8.0, 150 mM NaCl and 50 mM Tris pH 7.5, 150 mM NaCl buffer were mixed, respectively, 1: 1 with reservoir solution at room temperature. Data sets were collected at the Advanced Light Source, Berkley CA and processed with XDS (with highest resolution shell with a corresponding CC1/2 = ~ 0.1).^73^ Structures were determined by molecular replacement. Crystallographic and refinement statistics can be found in Supplementary Table 1. Final models were deposited to RCSB PDB with accession codes: 8EQ8 (14-3-3 β 130nY) and 8EQH (14-3-3 β 213nY). Details on crystallization, data collection and processing can be found in the Supplementary Materials.

### Pull down assays

Detailed methods for all pull down assays can be found in the Supplementary information provided.

### Mammalian Cell Culture

#### HEK293T cell culture and transfection

HEK293T cells were seeded at a density of 2.5×10^4^ cells per chamber in an 8-well Permanox® chamber slide (Thermofisher Scientific, catalog no. 177445) in a total volume of 400 μL. Cells were allowed to grow for 14 hours. Cells were then co-transfected using a total of 400 ng DNA per well with the pAcBac1-3-nitro-tyrosine (A7) machinery to encode 3-nitrotyrosine and a pAcBac1 expressing either 14-3-3β-eGFP WT, NLS-14-3-3β-eGFP WT, 14-3-3 β-eGFP K51E mutant, and 14-3-3β-eGFP with an amber TAG codon at positions 130 (130nY) or 213 (213nY) at a ratio of 1:8, as previously described.^35^ Cells were grown after transfection in the presence or absence of 300 μM 3-nitrotyrosine for 12-14 hours, after which the cells were prepared for immunofluorescence assays.

#### Fluorescence measurements of HEK293T 14-3-3-eGFP protein expressions

Replicates of the cell cultures described above were washed once with DPBS (ThermoFisher Scientific, catalog no. 14040133) and were detached from the 8-well chamber slides using 100 μL Trypsin-EDTA (ThermoFisher Scientific, catalog no. 25200056). Cells were then resuspended using 100 μL of DMEM (Corning, catalog no. 23-10-013-CV), and spun down into 1.7 mL Eppendorf tubes (VWR, catalog no. 87004-262). Cell pellets were washed using 500 μL DPBS, 2x by centrifugation and resuspension. The final cell pellets were resuspended in 100 μL DPBS and placed into a 96-well plate (ThermoFisher Scientific, catalog no. M33089); fluorescence measurements were taken using a BIOTEK® Synergy 2 Microplate Reader and data were plotted using GraphPad Prism.

#### Immunofluorescence of nitrated 14-3-3 β proteins in fixed HEK293T cells

Following transfection, cells were prefixed in each chamber with 200 μL of cell culture media and 200uL of 4% paraformaldehyde, 0.2% glutaraldehyde solution for 10 min on ice. Following prefixation, cells were washed with DPBS 3x for 5 min and incubated with 200 μL of fixation buffer (4% paraformaldehyde, 0.2% glutaraldehyde) for 30 minutes at room temperature. Cells are washed again with DPBS 3x for 5 min and incubated with 50mM glycine + 0.1% Triton-X in DPBS for 30 minutes. Cells were then incubated with Alexa Flour 647 Phalloidin (Thermofisher Scientific catalog no. A22287) in each chamber for 30 min at room temperature, followed by 3 washes with DPBS for 5 minutes. The resulting slide was mounted with a #1.5 cover slip (Thermofisher Scientific catalog no. 152250) using Prolong Gold Antifade with DAPI (Thermofisher Scientific catalog no. P36935). Each slide was imaged on a Zeiss LSM 780 NLO confocal microscope.

## ACKNOWLEDGEMENTS

We thank PA Karplus for guidance on manuscript preparation. We also thank HS Jang, E Van Fossen, CH Vesely, and AJ Eddins for scientific discussions. This work was supported in part by the GCE4All Biomedical Technology Development and Dissemination Center supported by National Institute of General Medical Science grant RM1-GM144227 [to R.A.M. and R.B.C.], National Institute of Health (NIH) 5R01GM131168-02 and 5R01GM1114653-04 [both to R.A.M], NIH Instrument Grant 1S10OD020111-01 [to the Oregon State University Mass Spectrometry Facility], the Medical Research Foundation at Oregon Health Sciences University [to R.B.C.], and the Collins Medical Trust [to R.B.C.]. We’d also like to acknowledge Beamline 5.0.1 and 5.0.2 of the Advanced Light Source, a U.S. DOE Office of Science User Facility under Contract No. DE-AC02-05CH11231, is supported in part by the ALS-ENABLE program funded by the National Institutes of Health, National Institute of General Medical Sciences, grant P30 GM124169-01. N.N.S. acknowledges that his work on 14-3-3 proteins was partially supported by the Russian Science Foundation (grant 19-74-10031).

## DATA ACCESIBILITY

The data that support the findings of this study are available from the corresponding author upon reasonable request. Structures and structure factors are available from RCSB PDB (https://www.rcsb.org) with accession numbers 8EQ8 and 8EQH.

