## Supplementary Material for "Genetic encoding of 3-nitro-tyrosine reveals the impacts of 14-3-3 nitration on client binding and dephosphorylation"

**SUPPLEMENARY METHODS**

**Protein Expression and Purification**

*14-3-3 β WT,* 130nY, *&* 213nY *Protein Expression & Purification*. Chemically competent BL21-ai cells were transformed with pDULE2-3nitroY-A7 machinery plasmid (3nY machinery plasmid)[1], and a pBAD expression plasmid harboring the 14-3-3 β gene with a C-terminally fused 6xHIS tag with either a codon encoding a Tyr residue, or an amber stop codon (TAG) at position 130 or 213 in its open reading frame. Transformed cells were grown in defined non-inducing media (NIM)[1,2] supplemented with 100 μg/ml ampicillin and 50 μg/ml spectinomycin overnight at 37ºC.[1] Then, 5 ml of the overnight NIM cultures were used to inoculate 500 mL of defined auto-induction media (AIM) containing the appropriate antibiotics. Cultures expressing protein with 3-nitro-tyrosine at positions 130 or 213 were supplemented with 0.5 mM 3-nitrotyrosine. Cell cultures were grown to an $OD_{600}$ of ~1.0-1.5 and were cold-shifted to 25ºC and allowed to express for 16-24 hours. Cells were resuspended in a lysis buffer containing 50 mM Tris pH 7.5, 500 mM NaCl, 20 mM imidazole pH 8, and 5% glycerol. Cells were lysed by microfluidization at 18,000 psi, and clarified lysate was obtained by centrifugation at 21,000 rcf for 30 minutes and at which point Ni-NTA resin was added to bind the $His_6$ tagged protein. The resin was washed with 30 resin bed volumes of lysis buffer, and then eluted with the lysis buffer supplemented with 300 mM imidazole. Elutions were desalted into 50 mM Tris, 150 mM NaCl pH 7.5 using with a HiPrep 26/10 Desalting column. Concentrations were determined using 14-3-3 β extinction coefficient $\varepsilon = 0.99$ mL $mg^{-1}$ $cm^{-1}$ . To remove the C-terminal $His_6$ tag, TEV protease was added at a 1:100 molar ratio of TEV to 14-3-3 β and allowed to incubate for 14-16 hours at 4 °C. After TEV protease cleavage, the protein mixtures were supplemented with 20 mM imidazole, and flowed through fresh Ni-NTA. The Flowthroughs contained pure 14-3-3 b, which was collected, concentrated to ~2 mL volume and injected onto HiLoad 26/600 Superdex 200 gel-filtration column (Cytiva, catalog no. 28989336). 14-3-3 b WT and 130nY were gel filtered into a buffer containing 50 mM HEPES pH 8, 150 mM NaCl. 14-3-3 213nY was gel filtered into a buffer containing 50 mM Tris pH 7.5, 150 mM NaCl. Fractions containing pure 14-3-3 b were collected and concentrated using Vivaspin 20, 10,000 MWCO (Cytiva, catalog no. 28932360) to a concentration of > 30 mg/mL, flash frozen in liquid $N_2$ and stored at -80°C.

*Expression and Purification of AVI-*bd*SUMOEu1-14-3-3 β Constructs.* Chemically competent BL21-ai cells were triply transformed with the pDULE2-3nitroY-A7 machinery plasmid, pBAD AVI-*bd*SUMOEu1 - 14-3-3 β (WT, 130TAG or 213TAG), and the pEVF-GST-BirA plasmid[3] which expresses a GST-BirA fusion protein to facilitate *in-vivo* biotinylation. Proteins were expressed identically as above for the 14-3-3 β constructs, except that the AIM was additionally supplemented with 25 μg/mL chloramphenicol for the pEVF-GST-BirA plasmid and 100 µM D-Biotin.[3,4] Proteins were purified by Ni-NTA as above, desalted into the same buffer, concentrated to ~ 200 µM, and flash-frozen in liquid $N_2$ for lon storage at -80º C.

*Expression and Purification of sfGFP-R18.* A pBAD-vector containing a genetic fusion encoding N-terminal 2 tandem 6xHIS tags, sfGFP, and a C-terminal R-18 peptide in its open reading frame was transformed into chemically competent BL21-ai cells. These cells were grown overnight in 2xYT containing 100 μg/mL ampicillin at 37ºC. 5 ml of this overnight culture was used to inoculate 500 ml of 2xYT containing the same concentration of ampicillin; cells were grown to an $OD_{600}$ of ~0.6 and expression was induced with 0.2% L-arabinose. Cells expressed overnight at 37ºC. Harvested cells were resuspended in 50 mM Tris pH 7.5, 500 mM NaCl, 5% glycerol, and 20 mM imidazole pH 8 (Buffer A). Cells were lysed by microfluidization, and crude cell extract was centrifuged at 21,000 rcf for 30-40 minutes. The supernatant was applied to 5 ml of Ni-NTA equilibrated in Buffer A and was rocked end over end for 30 minutes. The Ni-NTA was transferred to a column and washed with 15x bed volume of Buffer A, and then eluted with a similar buffer with 600 mM imidazole (Buffer B). Eluted Proteins were buffer exchanged using FPLC and a HiPrep 26/10 Desalting column into 50mM Tris pH 7.5 150 mM NaCl, and concentrated to ~3 mg/ml. Proteins were flash frozen in liquid $N_2$ and stored at -80°C.

*Expression and Purification of wild-type and S16-phosphorylated HSPB6.* Chemically competent B-95 Δ*A* Δ*fabR* Δ*serB* were transformed with pKW2-EFSep[5] and a pRBC plasmid[6] harboring a genetic fusion of an N-terminal 6xHis tag fused to *bd*SUMO[7] and HSPB6 (WT or S16TAG for directing pSer incorporation). Expressions were performed exactly as previously described.[6] Briefly, after transformation colonies containing pKW2-EFSep and the corresponding pRBC plasmid were grown overnight at 37 ºC on LB/agar plates containing 25 mg ampicillin and 7 mg/mL chloramphenicol. Approximately a dozen colonies were scraped and used to inoculate ZY-Non-inducing media (ZY-NIM) supplemented with 50 μg/ml ampicillin and 12.5 μg/ml chloramphenicol and grown at 37 ºC overnight.[6] The next day, 30 ml of overnight culture was used to inoculate 1.5 L of ZY-auto-inducing media (ZY-AIM) in a 2.8 L baffled Fernbach flask containing the same antibiotics, and cells were grown at 37 ºC until reach an $OD_{600}$ ~1.0-1.5, at which time the temperature was reduced to 18 ºC. After overnight expression, cells were harvested and resuspended with 50 mM Tris pH 7.5, 500 mM NaCl, 5% glycerol, 20 mM imidazole pH 8.0, 10 mM Sodium Fluoride (NaF), 5 mM Sodium Pyrophosphate (NaPPi), and 2 mM dithiothreitol (DTT) (Buffer C). Cells were lysed by microfluidization at 18,000 psi, and clarified lysate was obtained by centrifugation at 21,000 rcf for 30 minutes and at which point Ni-NTA resin was added to bind the $His_6$ tagged protein. The resin was washed with 30 resin bed volumes of lysis buffer, and then eluted with Buffer C supplemented with 300 mM imidazole. Proteins were then desalted with a HiPrep 26/10 column into 50 mM Tris pH 7.5, 150 mM NaCl, 5% glycerol, 20 mM imidazole, 10 mM NaF, 5 mM NaPPi, 2mM DTT. After desalting, 6xHIS-*bd*SUMO was cleaved with 30 nM 6xHIS-bdSenP1, on ice, for 45 minutes. The cleaved HSPB6 and pHSPB6 were purified by subtractive purification on Ni-NTA and concentrated to ~160 µM before flash-freezing in liquid $N_2$ and storage at -80$^o$ C.

*Expression and Purification of* bd*SENP1 and* bd*SENP1-Eu1*. *bd*SENP1 and *bd*SENP1-Eu1 proteases were expressed and purified as previously described.[7,8] Briefly, *bd*SENP1 and *bd*SENP1-Eu1 proteases were expressed from the pET28 vector in BL21ai cells in 2xYT media at 18$^o$C with 1 mM IPTG/0.1% (w/v) arabinose for 18 hrs. Cells were lysed in 50 mM Tris pH 7.5, 500 mM NaCl, 5% glycerol, 20 mM imidazole pH 8.0, and clarified lysate was obtained by centrifugation at 20,000 rcf for 30 min. Proteases were purified with Ni-NTA resin, eluted with 300 mM imidazole, desalted into 25 mM Tris, 300 mM NaCl, 10% glycerol, flash-frozen in liquid $N_2$ and stored at -80$^o$ C.

**Crystallization and Diffraction data collection**

*14-3-3 β 130nY*: Crystals were grown by the hanging drop method in which 14-3-3z 130nY protein at 20-30 mg/ml in 50 mM HEPES pH 8, 150 mM NaCl was mixed 1:1 with reservoir solution. The reservoir condition consisted of 20%-30% PEG 550 MME, 50 mM $MgCl_2$, and 100 mM HEPES pH 7-8.3. Crystals grew to full size (~200 x100 x 100 µm$^3$) in 3-5 day at room temperature.

*14-3-3 β 213nY*: Crystals were grown by the hanging drop method in which 14-3-3z 213nY protein at 20-30 mg/ml in 50 mM Tris pH 7.5, 150 mM NaCl was mixed 1:1 with reservoir solution. The reservoir condition consisted of 20%-27% PEG 3350, and 100 mM Bis-Tris pH 5.5-6.5. Supplementation with 50 mM $MgCl_2$ was optional and did not enhance or impede crystal growth. Crystals grew to full size (~250 x150 x 100 $\mu m^3$) in ~2 months.

All crystals were mounted onto cryo-loops (Hampton Research), soaked in artificial mother liquor containing reservoir with the addition of 25-30% glycerol and frozen in liquid $N_2$. Data sets were collected on beamlines 5.0.1 (for 14-3-3 b 130nY) and 5.0.2 (for 14-3-3 b 213nY) at the Advanced Light Source, Berkley CA, then individually processed and merged using XDS. Data cutoffs were determined by the highest resolution shell with a corresponding CC1/2 = ~ 0.1.[9] Structures were determined by molecular replacement using 2BQ0 as the search model. PHENIX and Coot were used to refine and build the models. Crystallographic and refinement statistics can be found in Supplementary Table 1. Final models were deposited to RCSB PDB with accession codes: 8EQ8 (14-3-3 β 130nY) and 8EQH (14-3-3 β 213nY).

**Pull down assays**

*sfGFP-R18 pull downs*. 0.6 mL total of Streptavidin-Sepharose (Cytiva, catalog no. 17511301) was equilibrated with 10 bed volumes of 50 mM Tris pH 7.5, 150 mM NaCl (Buffer E). The resin was divided equally into 3 tubes, and to each tube ~0.6 mg of either AVI-bdSUMOEu1-14-3-3 β WT, 130nY or 213nY was added. The biotinylated 14-3-3 b proteins were allowed to bind to Streptavidin-Sepharose for 25 minutes on ice, after which the resins were washed with 10 bed volumes of Buffer E. After washing, each 0.2 mL batch of resin containing a pre-loaded 14-3-3 b variant was split into 2 x 0.1 mL portions, and either ~1.5 mg of sfGFP-R18 or an equivalent volume of buffer E was added. Complexes were allowed to form for 30 minutes on ice. After incubating, the resin was washed 6 times: 3 times with a high-salt wash containing 50 mM Tris pH 7.5, 500 mM NaCl and 3 times with a low-salt wash containing 50 mM Tris pH 7.5, 150 mM

NaCl. The 14-3-3 b proteins (and any bound sfGFP-R18) were eluted by the addition of 300 nM *bd*SENP1-EuB protease in 50 mM Tris pH 7.5, 150 mM NaCl for 1 hr. Eluted protein concentrations were determined by A280 and loaded onto SDS-Page for analysis. Loads were normalized to concentrations of 14-3-3 eluted from the blank columns, representative of ~2 μg of 14-3-3 β.

*HSPB6 and pHSPB6 pull downs.* HSPB6 or pHSPB6 were first mixed with AVI-SUMOeu1-14-3-3 β WT, 130nY or 213nY at a molar ratio of 1 : 1.6 and allowed to complex for 45 min at 30 $^{o}$C. in 50 mM Tris pH 7.5, 150 mM NaCl and 2 mM DTT total concentration was added to prevent disulfide induced HSPB6 aggregation.[10] As a negative control, AVI-SUMOeu1-14-3-3 β WT was pre-cleaved with 500 nM *bd*SENP1-EuB for 35 minutes on ice and mixed with either HSPB6 or pHSPB6. During the incubation, 0.6 mL bed volume of Streptavidin-Sepharose was equilibrated with 10 bed volumes of 50 mM Tris pH 7.5, 150 mM NaCl, 10 mM NaF, 5 mM Sodium Pyrophosphate (NaPPi), 2mM DTT (Buffer F). After washing, 50 mL equilibrated Streptavidin-Sepharose of resin was added to each protein complex mixture and the biotinylated 14-3-3 b proteins were allowed to bind for 20 minutes. After binding, columns were further washed with 30 bed volumes of Buffer E, followed by elution with 300 nM *bd*SENP1-EuB. Eluted fractions were run on Phos-tag (Wako Chemicals, catalog no. AAL-107) and SDS-PAGE gels.

*HEK293T lysate pulldowns with site-specifically nitrated 14-3-3 β.* a pellet $5x10^8$ HEK293T cells were resuspended in 5 mL of 50 mM Tris pH 7.5, 150 mM NaCl, and 50 nM Calyculin-A , lysed by sonication and soluble lysate was obtained after centrifugation at 20,000 rcf for 20 min at 4$^{o}$C. 14-3-3 β proteins (wild-type, 130nY and 213nY) in 50 mM HEPES pH 8.5, 150 mM NaCl were conjugated to NHS-Sepharose according to manufacturer instruction, and a blank column was generated by substituting protein with 100 mM Tris. After conjugation, 200 µL of each resin was incubated with 1.2 mL of HEK293T lysate for 2 hours at room temperature. After incubation with lysate, the columns were washed with 50 bed volumes of high salt wash (50 mM Tris pH 7.5, 500 mM NaCl, 50 nM Calyculin-A) followed by 50 bed volumes of low salt wash (50 mM Tris pH 7.5, 150 mM NaCl, 50 nM Calyculin-A). The columns were then boiled in 1 bed volume of Laemli's buffer, and the supernatant was loaded on SDS-PAGE for analysis.

*Limited phosphatase hydrolysis of pHSPB6 in the presence of nitrated 14-3-3 β.* 50 µL reactions containing 8.3 µM pHSPB6 and 12.5 µM of 14-3-3 β WT, 130nY or 213nY in 50 mM Tris pH 7.5, 150 mM NaCl were incubated at 30 $^{o}$C with the addition of 1x PMP Buffer and the addition of 4 units of λ- phosphatase (New England Biolabs, catalog no. P0753S). Reactions were quenched at 0 (no phosphatase added), 5, 10, 15, 30, 60, and 180 minutes by acetone-precipitation. Precipitated proteins were pelleted and resuspended in Laemli's buffer for SDS-PAGE and Phos-tag gel analysis. A load equal to ~2 µg of pHSPB6 was loaded onto each lane of the gels corresponding to the samples time of quenching. Band densities were estimated using densitometry on Image J (Fiji) and normalized to the relative starting amount of pHSPB6.

**SUPPLEMENTARY TABLES**

**Supplementary Table 1.** Data Collection & Refinement Statistics

| | **14-3-3 β 130nY** | **14-3-3 β 213nY** |
|---|---|---|
| *Data collection* [a] | | |
| Space group | P 1 $2_1$ 1 | C 1 2 1 |
| Unit cell axes *a*, *b*, *c* (Å) | 52.40, 124.51, 55.23 | 85.29, 115.03, 79.49 |
| Resolution Limits (Å) | 47.68-1.50 (1.55-1.50) | 44.22-1.90 (1.95-1.90) |
| Unique Observations | 102662 (9538) | 52427 (3903) |
| Completeness | 99.8% (99.9%) | 99.7% (99.5%) |
| Multiplicity | 38.3 (27.8) | 26.1 (24.0) |
| Average I/ σ | 14.78 (0.44) | 11.75 (0.45) |
| $R_{meas}$ [b] (%) | 24.3 (699.9) | 16.5 (2261.0) |
| $CC^{1/2}$ | 99.9 (17.9) | 100.0 (19.7) |
| *Refinement* | | |
| $R_{cryst}$ / $R_{free}$ (%) | 17.7/20.4 | 18.6/21.7 |
| No. protein molecules | 2 | 2 |
| No. protein residues | 461 | 459 |
| No. water molecules | 399 | 144 |
| Total number atoms | 4069 | 3782 |
| rmsd bond angles (°) | 1.00 | 0.77 |
| rmsd bond lengths (Å) | 0.011 | 0.006 |
| <B> protein (Å$^2$) | 39.88 | 66.44 |
| <B> water (Å$^2$) | 47.70 | 64.53 |
| Ramachandran Plot (%) [c] | | |
| Favored | 98.45 | 98.43 |
| Outliers | 0.00 | 0.00 |
| PDB code | 8EQ8 | 8EQH |

[a] Numbers in parentheses correspond to values in the highest resolution bin
[b] $R_{meas}$ is the multiplicity-weighted merging *R*-factor[11]
[c] Ramachandran plot generated using Molprobity[12]

### SUPPLEMENTARY FIGURES

#### Supplementary Figure S1

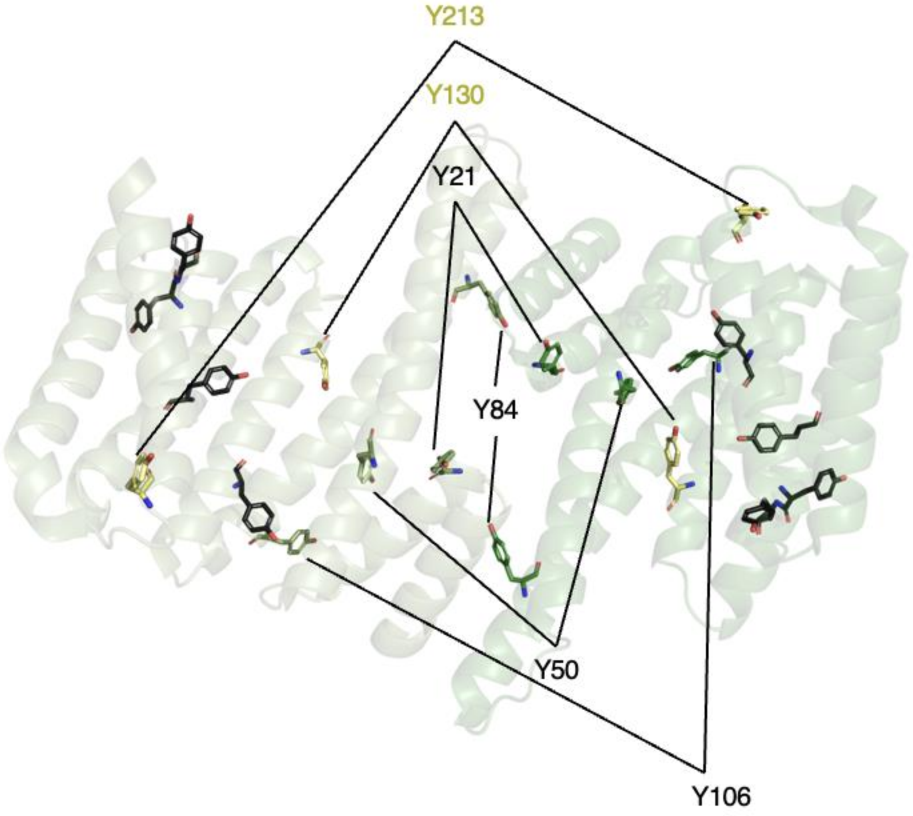

**Supplementary Figure S1.** Previously detected 3-nitrotyrosine residues in 14-3-3 proteins by proteomic mass spectrometry analyses,[13–18] mapped onto the 14-3-3 β structure (PDB: 2BQ0). Tyrosine residues not found to be nitrated are colored dark gray, those nitrated are shown in green, those nitrated that are discussed in this work are colored in yellow

**Supplementary Figure S2**

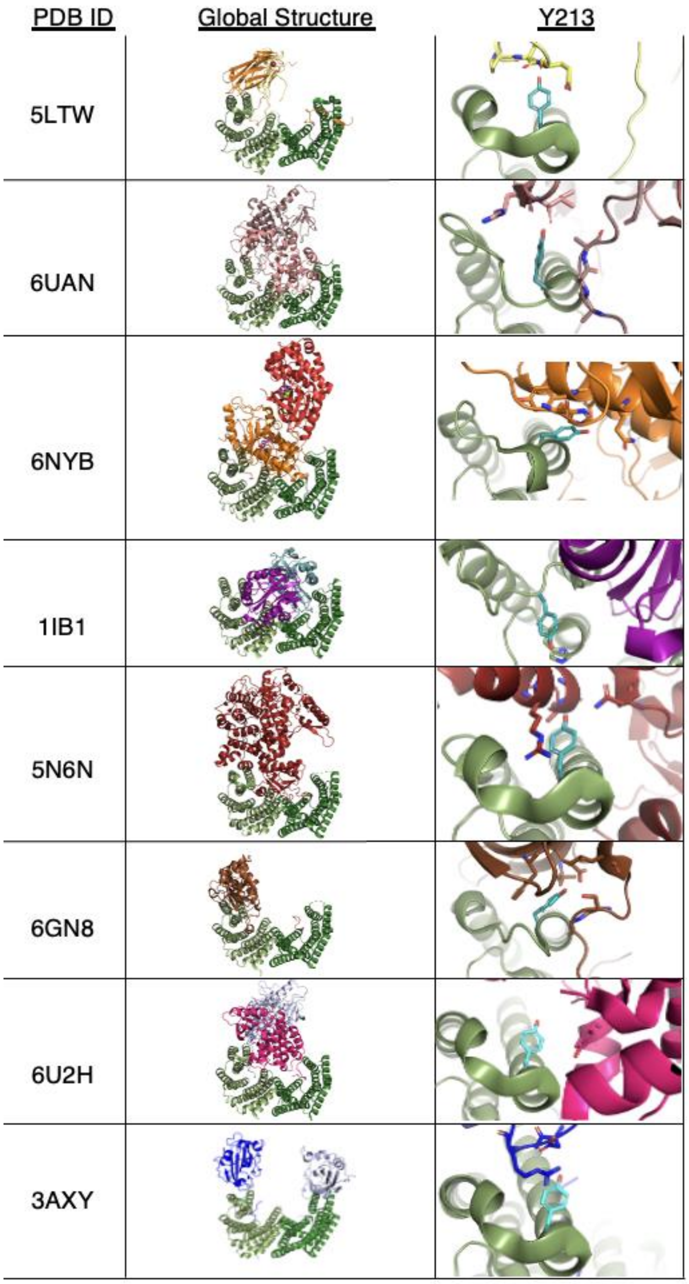

**Supplementary Figure S2.** Role of 14-3-3 Y213 (b numbering) within the so-called "hydrophobic patch" (Fig. 1 of main text) in various 14-3-3/client interactions. All 14-3-3 proteins are colored in green, with the 14-3-3 client-docking monomer in smudge-green and non-client docking monomer in forest-green. Right panels are a zoom-in of position Y213/client interactions, with relevant interacting client residues shown as sticks (various colors) with atoms 5 Å away; except for 1IB1, where 14-3-3 Y213 isn't client-interacting, but interacts with an intramolecular Histidine 166 (β-isoform numbering). PDB codes for each structure are listed in left column.

**Supplementary Figure S3**

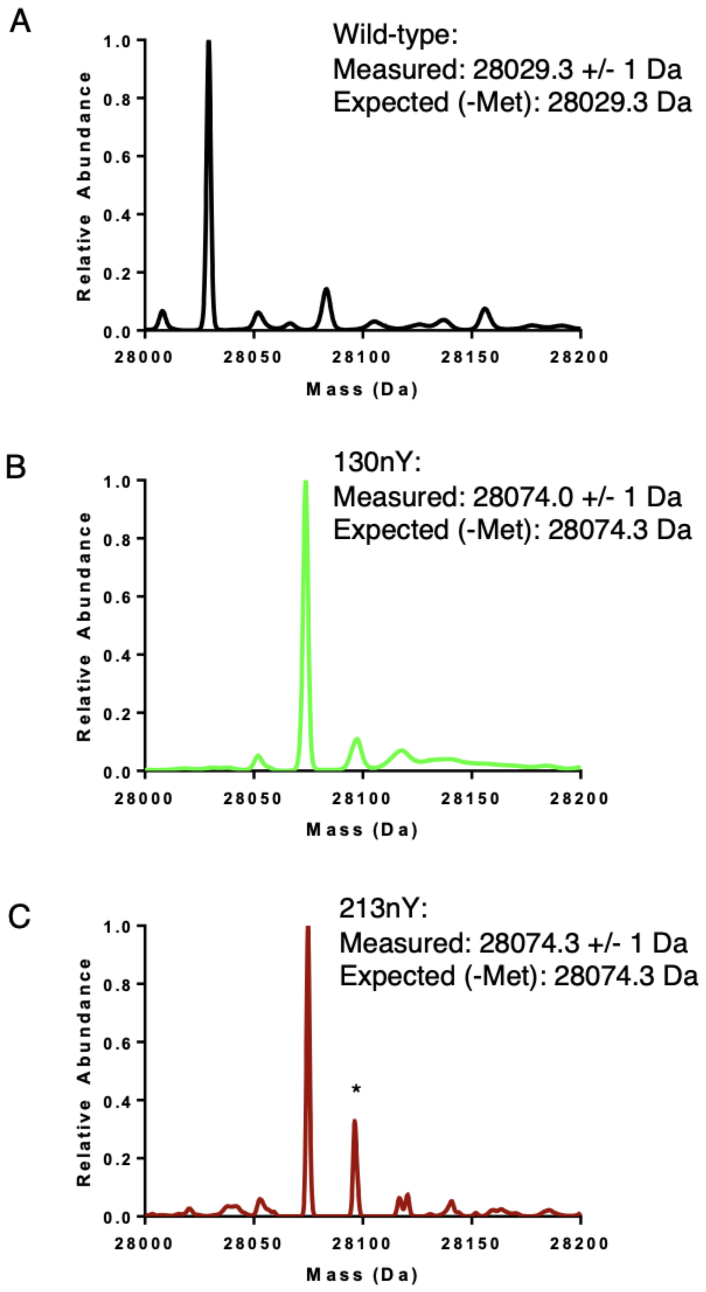

**Supplementary Figure S3.** Mass spectra of recombinantly expressed 14-3-3 β wildtype (A), 130nY (B) in and 213nY (C) in maroon line. * - Represents +22 Da $Na^{+}$ adduct of 213nY.

**Supplementary Figure S4**

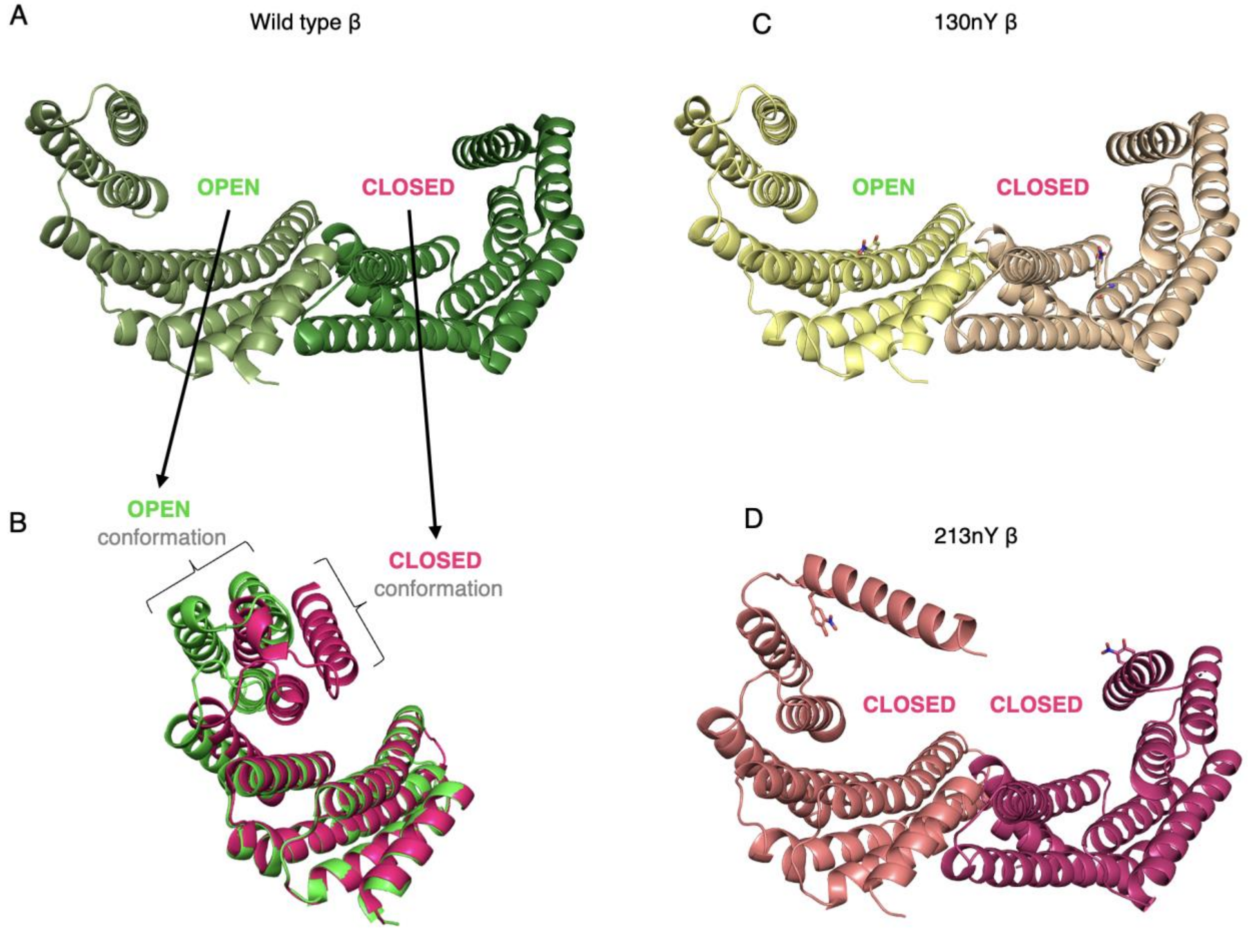

**Supplementary Figure S4.** Structural representation of the open/closed conformations of the 14-3-3 proteins. (A) Asymmetric biological dimer structure of apo 14-3-3 β showing the open (left) and closed (right) protomers (PDB: 2BQ0). (B) Overlay of the open (green) and closed (pink) conformations of apo 14-3-3 β (PDB: 2BQ0) highlighting relative positions of the C-terminal 3-helix bundle. (C) Structure of apo 14-3-3 β 130nY showing a similar asymmetric architecture as apo 14-3-3 β in panel A. (D) Structure of apo 14-3-3 β 213nY reveals a closed/closed architecture more commonly seen in phospho-peptide bound 14-3-3 structures.

**Supplementary Figure S5**

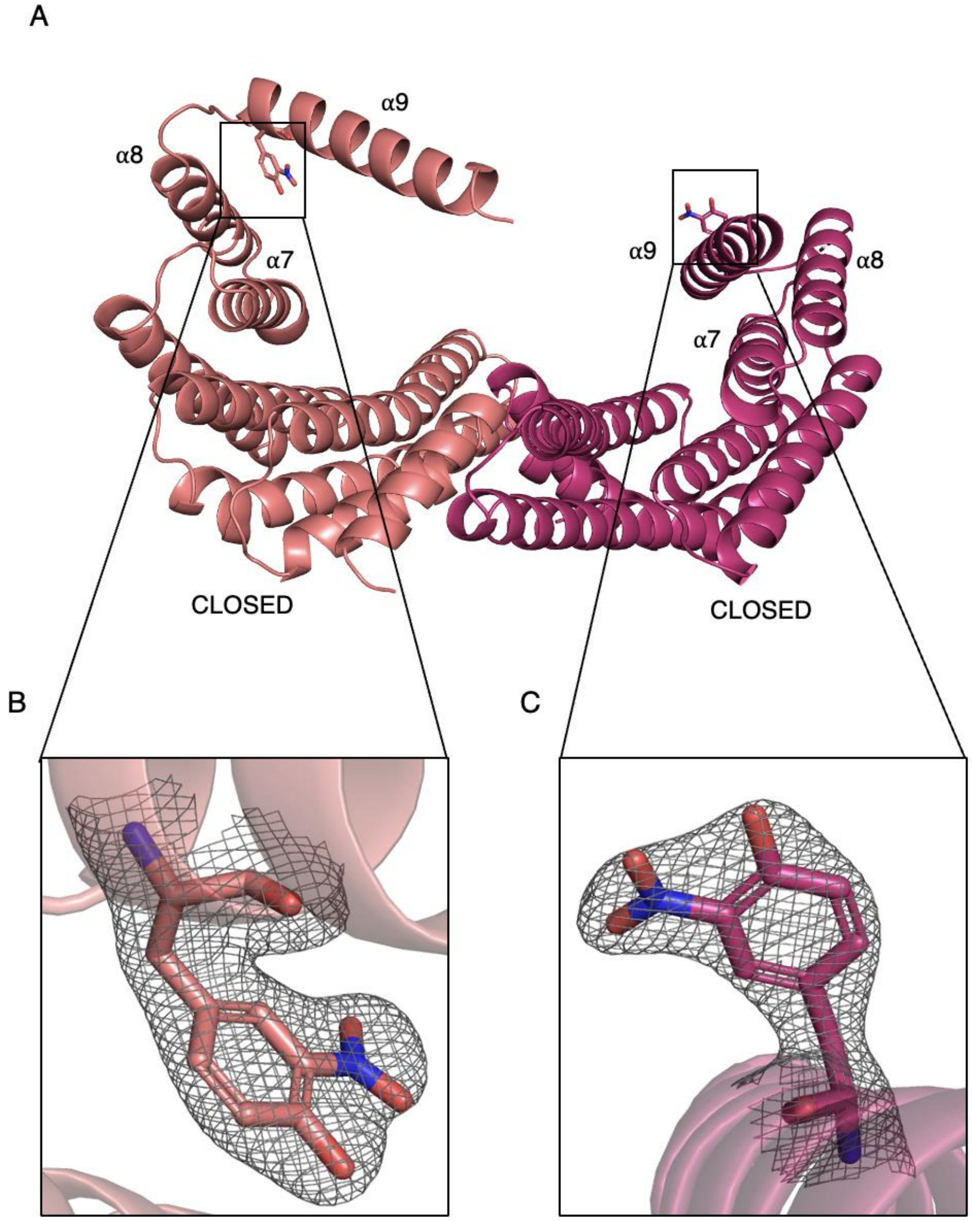

**Supplementary Figure S5.** Overall structure of 14-3-3 β 213nY. (A) 14-3-3 β 213nY crystalized with both protomers in the closed conformation where the terminal helix (a9) of one protomer (pink) has disengaged with its three-helix bundle and swung outward to connect and domain-swap with a neighboring 14-3-3 molecule in the crystal lattice (not shown, see Supplementary Figure S6). The other protomer (magenta) is not domain swapped and adopts a closed conformation. (B) Electron-density ($2F_o$-$F_c$, 1σ) of the 3-nitro-tyrosine moiety at position Y213 for the domain-swapped protomer and the non-domain swapped protomers.

**Supplementary Figure S6**

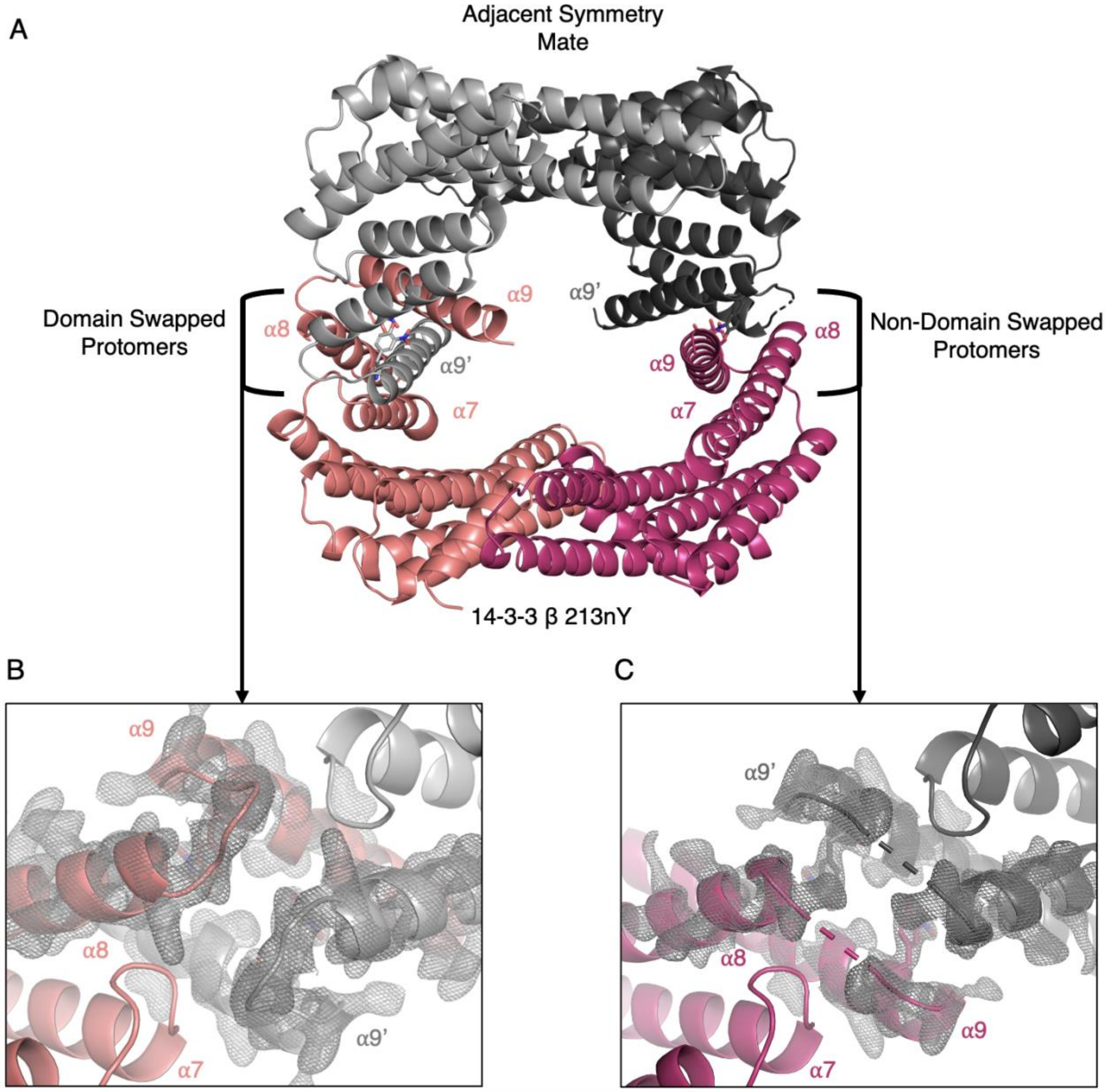

**Supplementary Figure S6.** Domain swap in the 14-3-3 β 213nY structure. (A) Global structure of the 14-3-3 β 213nY biological dimer (pink, magenta) with an adjacent crystallographic symmetry mate (light grey, dark grey) with a-helices labelled shows how a9 and a9' have swapped to engage with their neighboring 14-3-3 molecule rather than with the from which is originates. (B) Electron density ($2F_o$-$F_c$, 1σ) of the loop bridging helices a8 and a9 showing clear density for the domain-swapped protomer (pink) with its crystallographic symmetry mate (light grey). (C) Electron density ($2F_o$-$F_c$, 1σ) of the loop bridging helices a8 and a9 showing the absence of density within the bridging loop. The protomer of interest alpha helices are labeled α7, 8 and 9, respectively and alpha helix 9 of the adjacent symmetry mate is labeled α9'.

**Supplementary Figure S7**

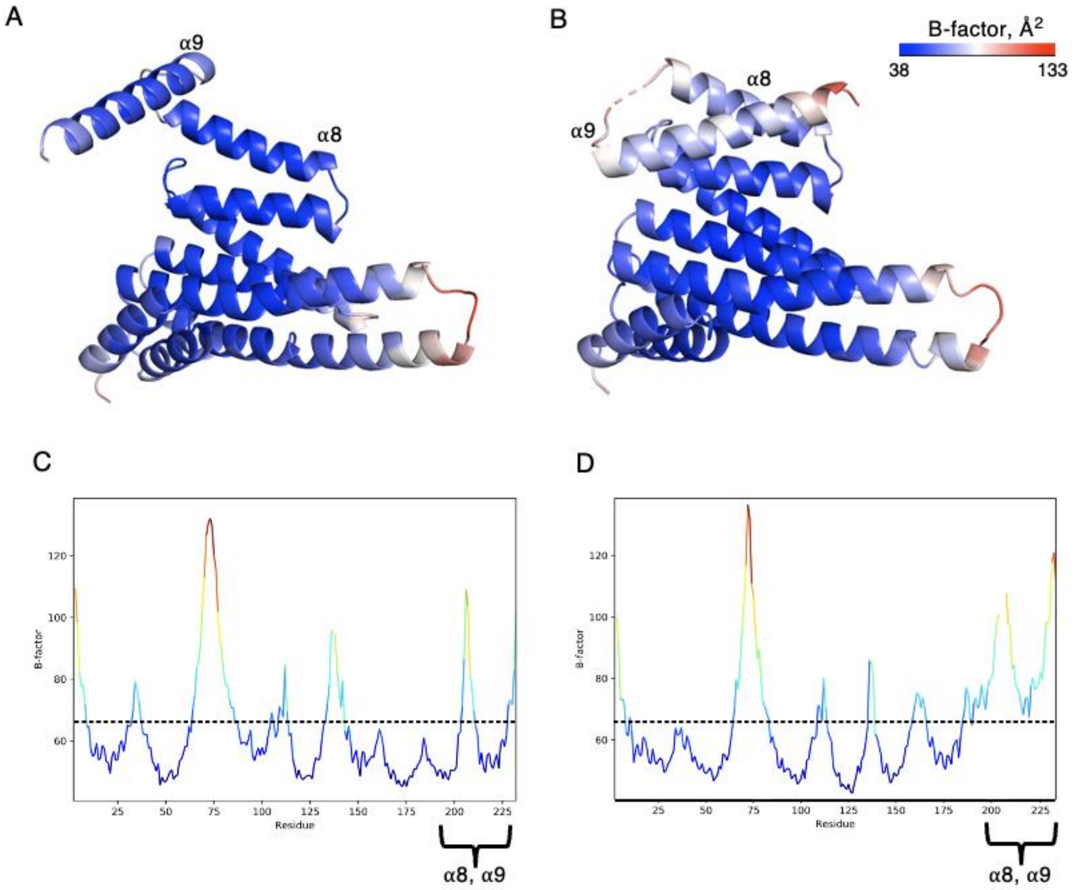

**Supplementary Figure S7.** B-factor analysis of the domain swapped 14-3-3 β 213nY structure. (A) The structures of the domain-swapped protomer and (B) the non-domain swapped protomer colored by B-factor from blue to white to red with scale bar shown to the right; the a-helices are labeled above each helix. (C) Plots depicting the average B-factor of each residue for the domain-swapped protomer and (D) the non-domain swapped protomer. Dashed-line represents the overall B-factor of each protomer. The region spanning helices α8 and α9 are labelled under the graph for each protomer.

**Supplementary Figure S8**

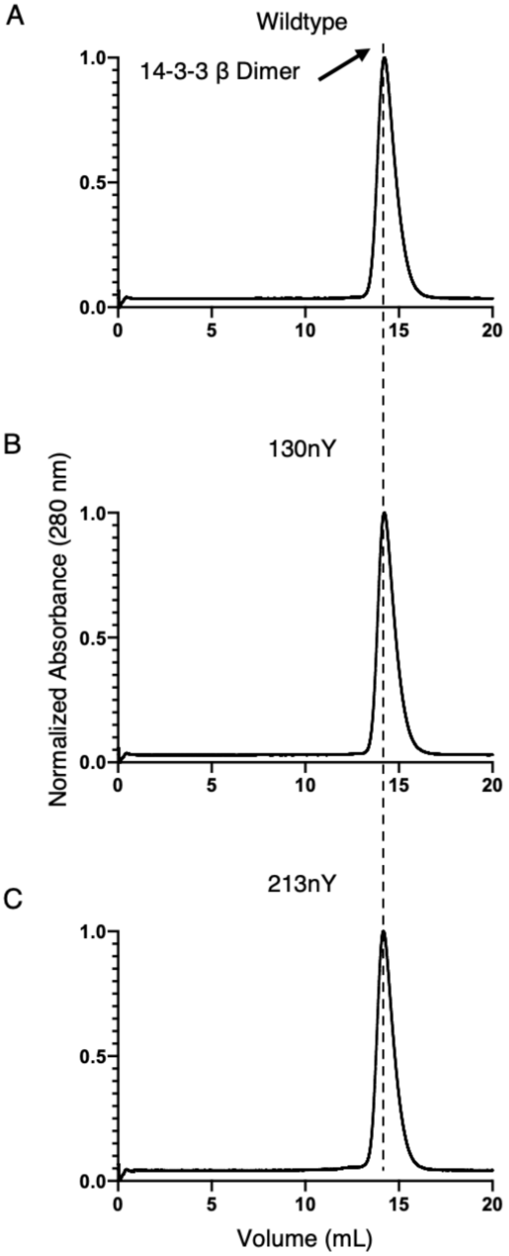

**Supplementary Figure S8.** Analytical size-exclusion chromatography of the 14-3-3 β wild-type (top) 130nY (middle) and 213nY (C) proteins. The dotted line represents the elution volume of the 14-3-3 dimer, at 14.2 mL. The identical elution times are indicative that all variants adopt the same oligomeric state (dimeric).

**Supplemental Figure S9**

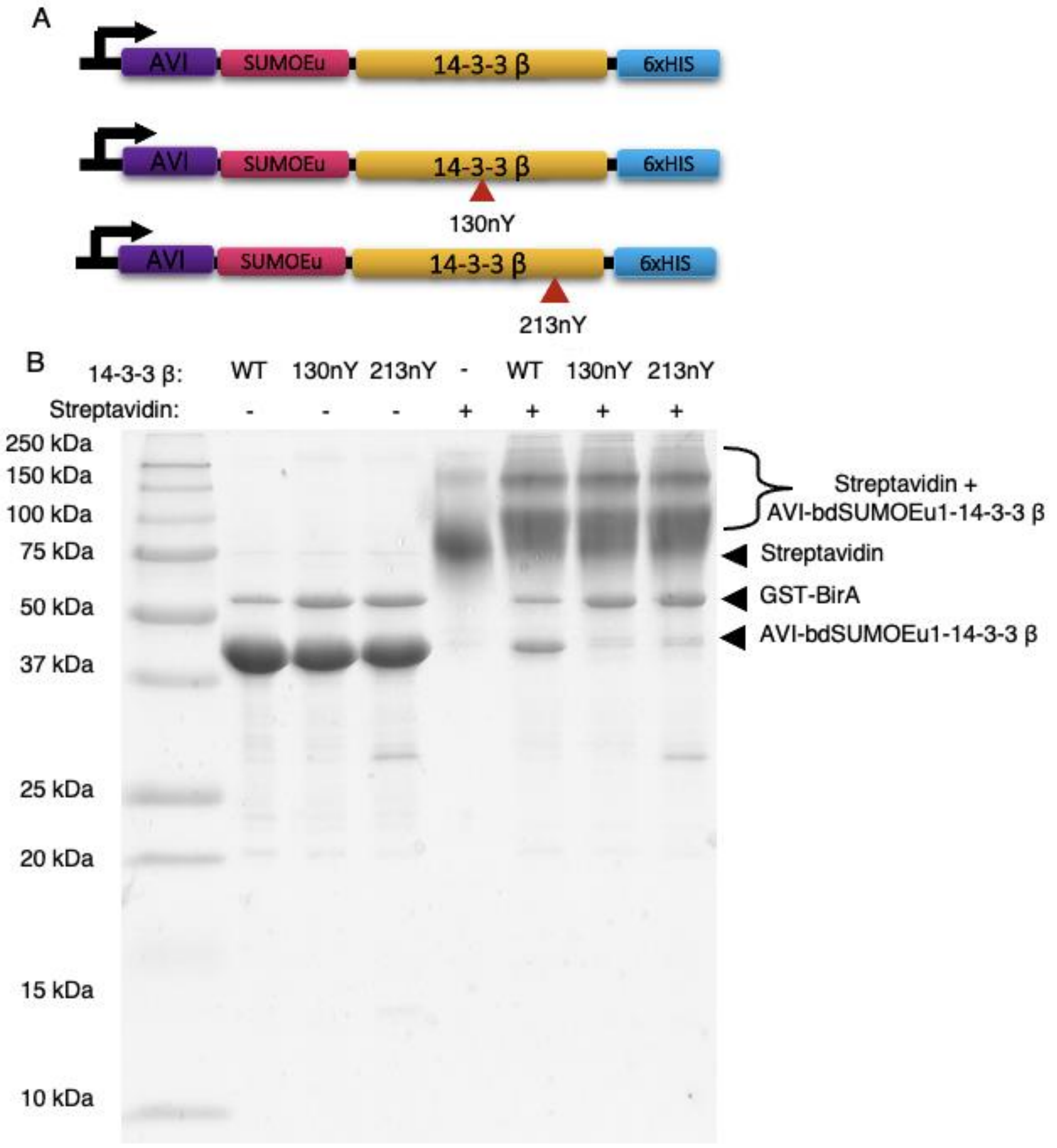

**Supplementary Figure S9.** Construction and synthesis of AVI-SUMO-tagged, nitrated 14-3-3 proteins. (A) 14-3-3 fusion protein constructs used in sfGFP-R18 and pHSPB6 pull down assays contained an N-terminal AVI tag for *in vivo* biotinylation, fused to a SUMO protein (bdSUMOEu1) followed by 14-3-3 containing either a Tyr triplet codon at positions Y130 and Y213 or an amber (TAG) at positions Y130 or Y213. (B) SDS-Page mobility shift assay demonstrating successful biotinylation of ~90% of 14-3-3 wild-type (WT) and near-complete biotinylated of 130nY and 213nY; Streptavidin was added to the 14-3-3 fusion proteins after boiling in Laemli's buffer for 5 mins.

**Supplementary Figure S10**

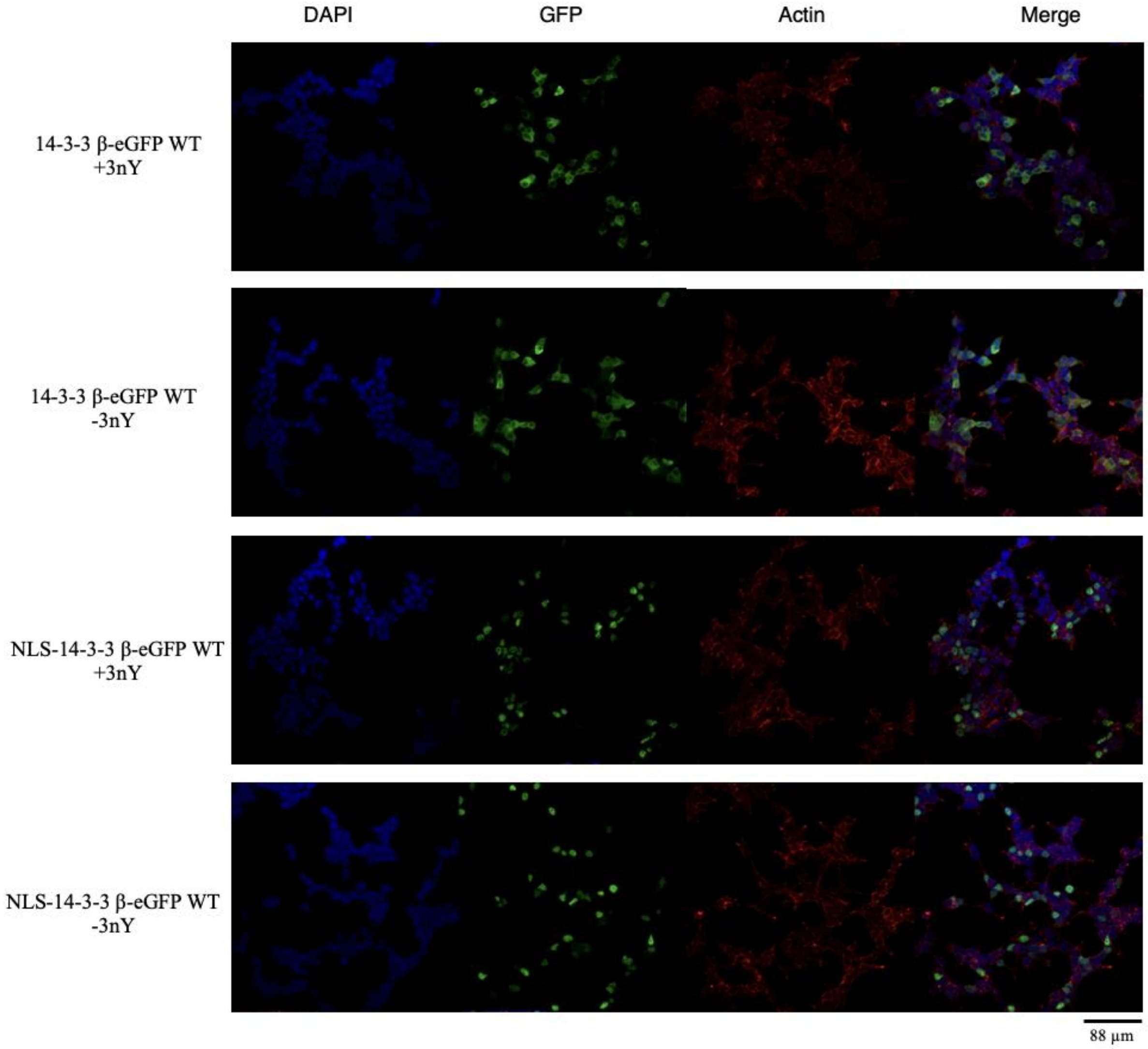

### Supplementary Figure S10 cont.

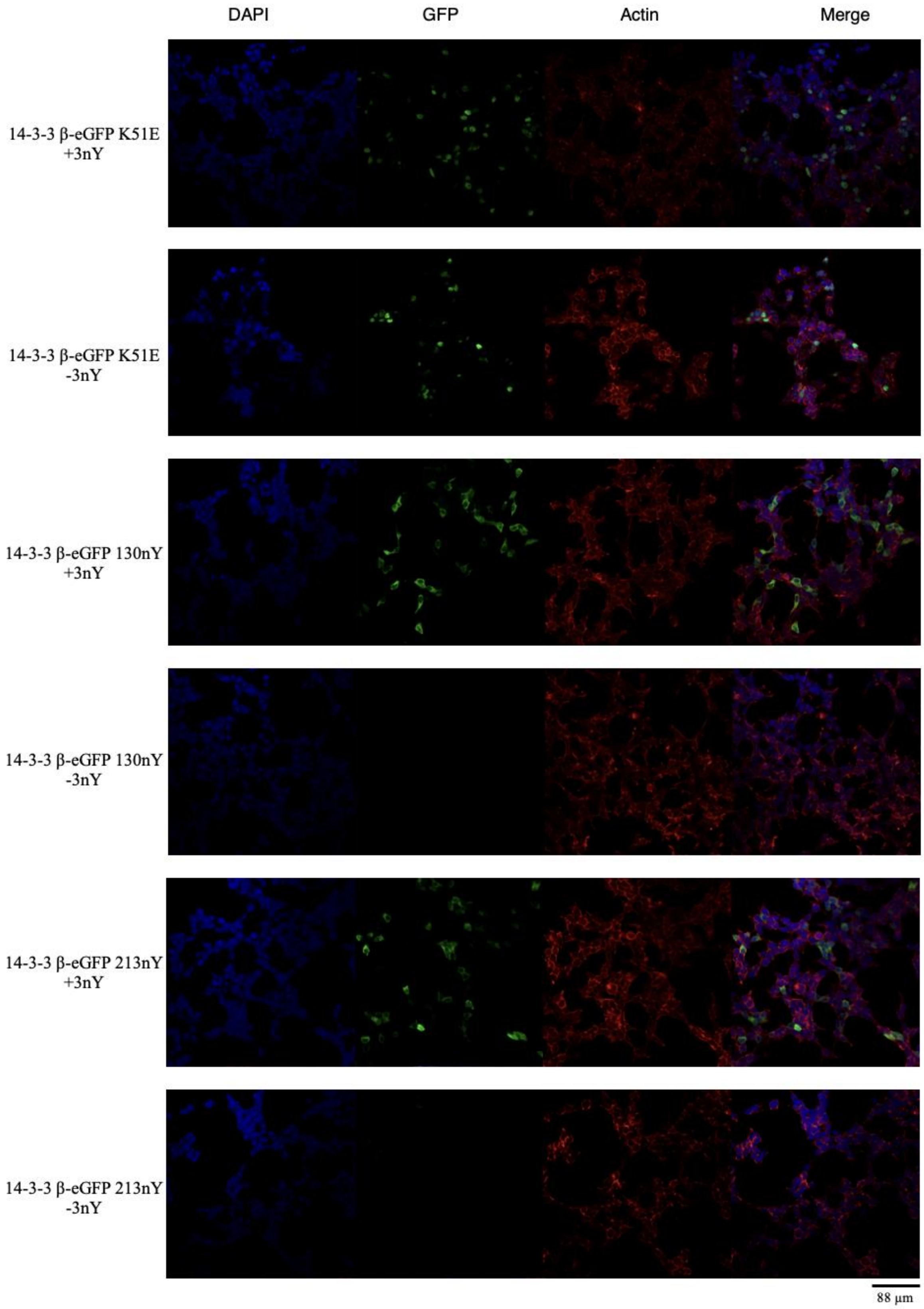

**Supplementary Figure S10.** Unmerged channels of DAPI (blue) channel, GFP (green) channel, and Actin (red) channel, and a merged image of all channels for the expression of 14-3-3 β – eGFP fusion proteins containing a wildtype (WT) sequence or containing an N-terminal nuclear localization sequence (NLS), a K51E mutation, TAG interrupted gene at position Y130 (130nY), or TAG interrupted gene at position Y213 (213nY). Images were taken from cultures with 0.3 mM 3nY (+3nY), or without (-3nY). Black scale bar, lower right represents 88 µm.

**Supplementary Figure S11**

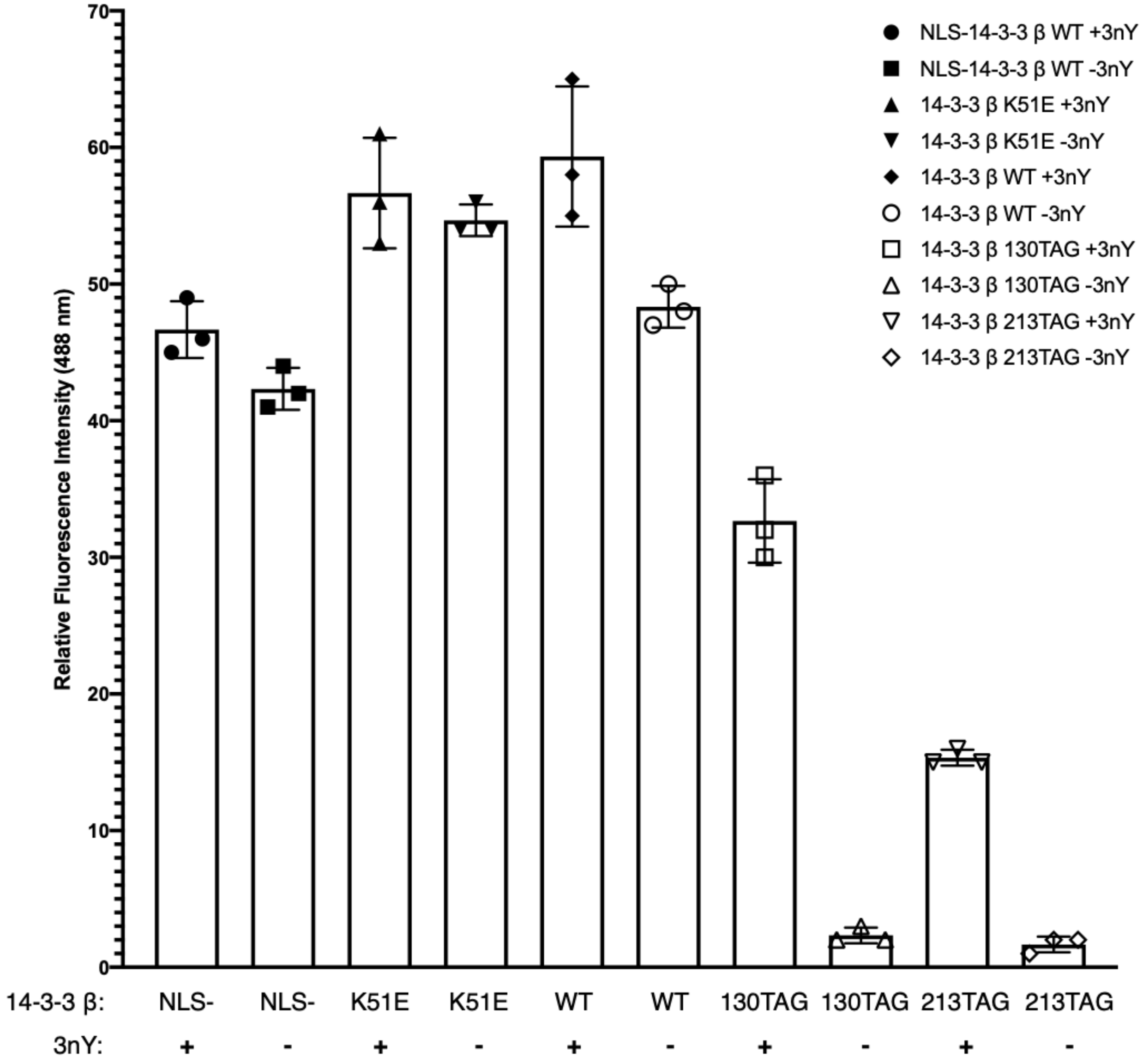

**Supplementary Figure S11.** Relative fluorescence intensities of transiently transfected HEK293T cells containing the indicated 14-3-3 β-eGFP fusion constructs and with (+3nY) 0.3 mM 3-nitrotyrosine in the media, or without (-3nY) 3-nitrotyrosine supplementation.

**Supplementary Figure S12**

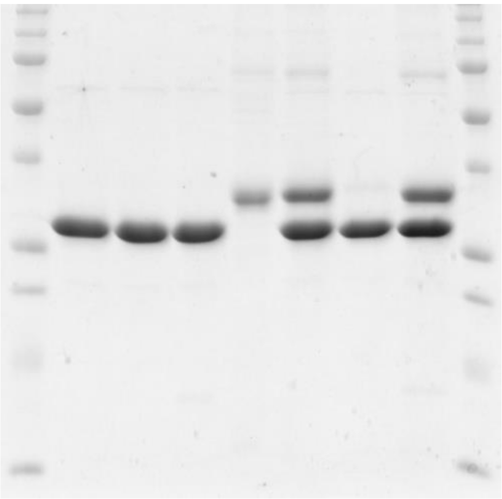

**Supplementary Figure S12.** Gel used for cropped image in Figure 3B.

**Supplementary Figure S13**

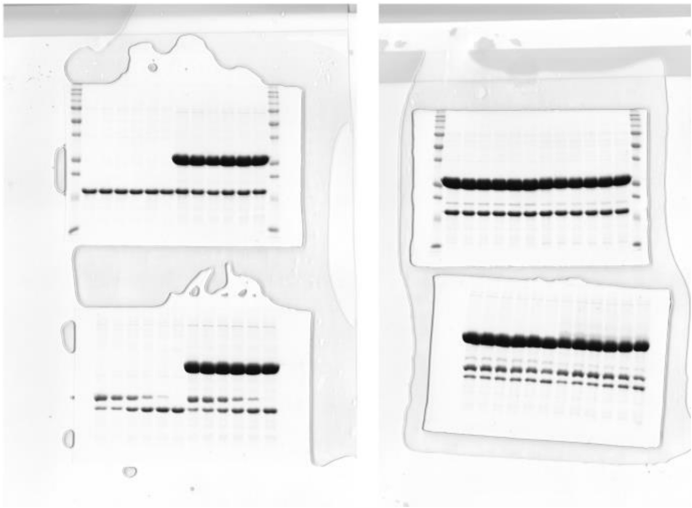

**Supplementary Figure S13.** Gels used for cropped gel image in Figure 6.

**PROTEIN SEQUENCES**

***E. coli expression sequences***

**>14-3-3 b - TEV – His$_6$**
MGMDKSELVQKAKLAEQAERYDDMAAAMKAVTEQGHELSNEERNLLSVAYKNVVG
ARRSSWRVISSIEQKTERNEKKQQMGKEYREKIEAELQDICNDVLELLDKYLIPNATQPE
SKVFYLKMKGDYFRYLSEVASGDNKQTTVSNSQQAYQEAFEISKKEMQPTHPIRLGLAL
NFSVFYYEILNSPEKACSLAKTAFDEAIAELDTLNEESYKDSTLIMQLLRDNLTLWTSEN
QGDEGENLYFQSGTHHHHHH
Note: Residues underlined = sites of nitroY incorporation

**>AVI-SUMOEu1-14-3-3 b - TEV-His$_6$**
MGLNDIFEAQKIEWHEGSGGGAHINLKVKGQDGNEVFFRIKRSTQLKKLMNAYCDRQS
VDMKAIAFLFKGRRLRAERTPDELEMEDGDEIDAMLHQTGGSGMGMDKSELVQKAKL
AEQAERYDDMAAAMKAVTEQGHELSNEERNLLSVAYKNVVGARRSSWRVISSIEQKTE
RNEKKQQMGKEYREKIEAELQDICNDVLELLDKYLIPNATQPESKVFYLKMKGDYFRY
LSEVASGDNKQTTVSNSQQAYQEAFEISKKEMQPTHPIRLGLALNFSVFYYEILNSPEKA
CSLAKTAFDEAIAELDTLNEESYKDSTLIMQLLRDNLTLWTSENQGDEGENLYFQSGTH
HHHHH

**>His$_6$- *bd*SUMO-HSPB6**
MGSSHHHHHHSGSAAGGEEDKKPAGGEGGGAHINLKVKGQDGNEVFFRIKRSTQLKKL
MNAYCDRQSVDMTAIAFLFDGRRLRAEQTPDELEMEDGDEIDAMLHQTGGSGGEIPVP
VQPSWLRRASAPLPGLSAPGRLFDQRFGEGLLEAELAALCPTTLAPYYLRAPSVALPVA
QVPTDPGHFSVLLDVKHFSPEEIAVKVVGEHVEVHARHEERPDEHGFVAREFHRRYRLP
PGVDPAAVTSALSPEGVLSIQAAPASAQA
Note:  Residues underlined = site of pSer incorporation

**>His$_{12}$- sfGFP-R18**
MGHHHHHHGSTGTGSGSSGHHHHHHGGSSKGEELFTGVVPILVELDGDVNGHKFSVRG
EGEGDATNGKLTLKFICTTGKLPVPWPTLVTTLTYGVQCFSRYPDHMKRHDFFKSAMPE
GYVQERTISFKDDGTYKTRAEVKFEGDTLVNRIELKGIDFKEDGNILGHKLEYNFNSHN
VYITADKQKNGIKANFKIRHNVEDGSVQLADHYQQNTPIGDGPVLLPDNHYLSTQSKLS
KDPNEKRDHMVLLEFVTAAGITHGMDELYKGSGSTSGGTGPHCVPRDLSWLDLEANM
CLP

***HEK293T expression sequences***

**> 14-3-3 β-eGFP**
MGMDKSELVQKAKLAEQAERYDDMAAAMKAVTEQGHELSNEERNLLSVAYKNVVG
ARRSSWRVISSIEQKTERNEKKQQMGKEYREKIEAELQDICNDVLELLDKYLIPNATQPE

SKVFYLKMKGDYFRYLSEVASGDNKQTTVSNSQQAYQEAFEISKKEMQPTHPIRLGLAL
NFSVFYYEILNSPEKACSLAKTAFDEAIAELDTLNEESYKDSTLIMQLLRDNLTLWTSEN
QGDEGGGGSGGGSVSKGEELFTGVVPILVELDGDVNGHKFSVSGEGEGDATYGKLTLK
FICTTGKLPVPWPTLVTTLTYGVQCFSRYPDHMKQHDFFKSAMPEGYVQERTIFFKDDG
NYKTRAEVKFEGDTLVNRIELKGIDFKEDGNILGHKLEYNYNSHNVYIMADKQKNGIKV
NFKIRHNIEDGSVQLADHYQQNTPIGDGPVLLPDNHYLSTQSKLSKDPNEKRDHMVLLE
FVTAAGITLGMDELYKWSHPQFEKGGGSGGGSGGSAWSHPQFEKGKPIPNPLLGLDSTH
HHHHH

Note: Residues underlined = sites of nitroY incorporation

> **14-3-3 β-eGFP K51E**

MGMDKSELVQKAKLAEQAERYDDMAAAMKAVTEQGHELSNEERNLLSVAY**E**NVVGA
RRSSWRVISSIEQKTERNEKKQQMGKEYREKIEAELQDICNDVLELLDKYLIPNATQPES
KVFYLKMKGDYFRYLSEVASGDNKQTTVSNSQQAYQEAFEISKKEMQPTHPIRLGLAL
NFSVFYYEILNSPEKACSLAKTAFDEAIAELDTLNEESYKDSTLIMQLLRDNLTLWTSEN
QGDEGGGGSGGGSVSKGEELFTGVVPILVELDGDVNGHKFSVSGEGEGDATYGKLTLK
FICTTGKLPVPWPTLVTTLTYGVQCFSRYPDHMKQHDFFKSAMPEGYVQERTIFFKDDG
NYKTRAEVKFEGDTLVNRIELKGIDFKEDGNILGHKLEYNYNSHNVYIMADKQKNGIKV
NFKIRHNIEDGSVQLADHYQQNTPIGDGPVLLPDNHYLSTQSKLSKDPNEKRDHMVLLE
FVTAAGITLGMDELYKWSHPQFEKGGGSGGGSGGSAWSHPQFEKGKPIPNPLLGLDSTH
HHHHH

Note: Bolded E is the site of K → E mutation

> **NLS-14-3-3 β-eGFP**

MPAAKRVKLDGGSGGMDKSELVQKAKLAEQAERYDDMAAAMKAVTEQGHELSNEER
NLLSVAY**K**NVVGARRSSWRVISSIEQKTERNEKKQQMGKEYREKIEAELQDICNDVLEL
LDKYLIPNATQPESKVFYLKMKGDYFRYLSEVASGDNKQTTVSNSQQAYQEAFEISKKE
MQPTHPIRLGLALNFSVFYYEILNSPEKACSLAKTAFDEAIAELDTLNEESYKDSTLIMQL
LRDNLTLWTSENQGDEGGGGSGGGSVSKGEELFTGVVPILVELDGDVNGHKFSVSGEG
EGDATYGKLTLKFICTTGKLPVPWPTLVTTLTYGVQCFSRYPDHMKQHDFFKSAMPEG
YVQERTIFFKDDGNYKTRAEVKFEGDTLVNRIELKGIDFKEDGNILGHKLEYNYNSHNV
YIMADKQKNGIKVNFKIRHNIEDGSVQLADHYQQNTPIGDGPVLLPDNHYLSTQSKLSK
DPNEKRDHMVLLEFVTAAGITLGMDELYKWSHPQFEKGGGSGGGSGGSAWSHPQFEK
GKPIPNPLLGLDSTHHHHHH
